# Inversions support both parallel and location-specific adaptations in snail ecotypes

**DOI:** 10.64898/2026.01.05.697708

**Authors:** Basile Pajot, Thomas Broquet, Le Qin Choo, Pierre Barry, Rui Faria, Roger Butlin, Kerstin Johannesson, Alan Le Moan

## Abstract

Replicated hybrid zones between ecotypes established over within shore environmental gradients provide an opportunity to study the genomic architecture of barriers to gene flow. The marine snail *Littorina fabalis* segregates locally into dwarf and large ecotypes over wave-exposure gradients on northwestern European shores. Previous work focusing on a hybrid zone in Sweden revealed strong genetic differentiation between the ecotypes, concentrated in 12 putative chromosomal inversions. Here we compare the Swedish hybrid zone with samples from France distributed across a similar wave-exposed gradient. Our aims were to test if similar exposure gradients promote parallel ecotype distributions and hybrid zones, and if similar genomic architectures contribute to divergence and adaptation across the gradients. Unpredictably, we found that the shell size cline was reversed in France compared to Sweden, with small individuals occupying the more-sheltered end of the environmental gradient in Sweden but the more-exposed end in France. We also observed a cline in shell colour in France, whereas nearly all Swedish snails were yellow. Using whole-genome sequencing, we found similar levels of genetic differentiation between ecotypes in both places. Most of the differences were accounted for by the same 15 inversions, and the arrangement clines showed similar associations to the wave-exposure gradient in both hybrid zones. These inversions were enriched in SNPs differentiating the ecotypes that were either specific to one hybrid zone or showed reversed cline patterns between zones. Genome-wide association studies (GWAS) detected significant associations between genomic regions within inversions and shell size in Sweden, while one inversion was associated with colour in France. Our results show that the same inversions play a dual role: they support ecotype differences across similar environmental gradients in distant locations, while also contribute site-specific variation contributing to local adaptation.

## Introduction

The evolution of ecotypes that occur along similar environmental gradients in different geographic regions provides valuable systems to study the genomic basis of adaptive divergence and establishment of reproductive barriers (e.g. Jones et al., 2012; Morales et al., 2019). By comparing genomic landscapes across replicate ecotype hybrid zones located in distinct geographical sites, we can distinguish shared genomic architectures, that promote local adaptation and establish barriers to gene flow, from site-specific genetic differences (Pal et al., 2025). Such comparisons have earlier revealed that genomic architectures contributing to ecotype divergence frequently involve chromosomal inversions (Harringmeyer and Hoekstra, 2022; Huang et al., 2020; Jones et al., 2012; Meyer et al., 2025, 2024; Nicolas et al., 2025), which maintain reproductive isolation, sometimes across remarkably small spatial scales (Le Moan et al., 2024; Westram et al., 2021).

Determining the nature of the barrier generated by these inversions remains challenging because reproductive isolation often results from a combination of diverse processes involving not only local adaptation, but also intrinsic genetic incompatibilities and assortative mating (Butlin and Smadja, 2018; Johannesson et al., 2024). The reduced recombination between inversion arrangements can, furthermore, generate strong associations between multiple barrier types (Wellenreuther and Bernatchez, 2018). In addition, many ecotype-forming species have acquired such multifaceted barriers during an allopatric phase of divergence that preceded secondary contact (Le Moan et al., 2016; Rougemont et al., 2017; Rougeux et al., 2019). Under this scenario, replicated associations of ecotypes with similar environmental gradients may reflect the spatial spread of divergent lineages rather than parallel adaptation (Bierne et al., 2013). Consequently, associations between genetic differentiation and environmental gradients do not necessarily imply direct evidence for environmental selection acting on those inversions (Bierne et al., 2011; Ravinet et al., 2017).

Two complementary approaches relying exclusively on observations from natural populations can help to resolve the nature of barriers acting in these inversion-structured systems. First, when allopatrically diverged ecotypes come into secondary contact in multiple locations, intrinsic barrier loci may associate with opposing ends of environmental gradients depending on historical contingency (Bierne et al., 2011). Such reversed clines would manifest as follows: if an inversion contains loci causing genetic incompatibilities (but no locally-adaptive loci), historical events would determine which arrangement colonizes which habitat in each location. Thus, arrangement A might associate with one end of the environmental gradient at one site but with the opposite end at another, despite identical environmental conditions. In contrast, if the inversion harbours adaptive loci, arrangement A should consistently associate with the same environmental conditions across all sites. While theoretically predicted, this signature of intrinsic barriers remains to be empirically tested across replicate contact zones (but see Riginos and Cunningham, 2005). Second, association mapping in natural populations (e.g., Genome-Wide Association Study, GWAS) can be used to link inversions to barrier-related phenotypes, thereby revealing the selective forces maintaining them (Hager et al., 2022; Koch et al., 2022). Despite this potential, few studies have established explicit phenotypic links between inversions and specific barrier mechanisms in natural systems (e.g., Harringmeyer and Hoekstra, 2022; Jones et al., 2012; Koch et al., 2022, 2021).

Chromosomal inversions not only shape patterns of parallel differentiation between ecotypes but may also affect site-specific evolutionary responses (Faria et al., 2019). At a given geographical site, demographic history and local bottlenecks may reduce neutral diversity, particularly within polymorphic inversion arrangements, which have lower effective population sizes than collinear genomic regions (Berdan et al., 2023). These demographic signatures persist longer in inversions due to suppressed recombination, similar to patterns near centromeres (Duranton et al., 2018). Inversions can also respond to site-specific environmental pressures, though their role in local adaptation may be complex. On one hand, inversions may facilitate adaptation: locally advantageous mutations within inversions already associated with ecotype divergence are protected from gene flow, making them more likely to spread than mutations in collinear regions (Faria and Navarro, 2010; Lenormand, 2002). This process may enable location-specific adaptation after an inversion becomes established in a particular region. On the other hand, inversions may constrain the adaptive potential when spatial variation in ecological conditions alters the selective value of particular associations among adaptive loci (Roesti et al., 2022). Thus, while inversions can clearly contribute to parallel genetic differences along well-replicated environmental gradients (Westram et al., 2022), their impact becomes less predictable when gradients differ ecologically across sites or when allopatric divergence contributes to the accumulation of intrinsic barrier loci within inversions.

The marine snails of the genus *Littorina* have been developed into useful models to study speciation and the consequences of chromosomal inversions for local adaptation (Johannesson et al., 2024; Reeve et al., 2024). The genus includes several species forming ecotypes that differ in phenotypic traits such as shell morphology, size, behaviour, and physiology (Reid, 1996; Johannesson, 2003; Johannesson et al., 2010) across ecological gradients (Morales et al., 2019; Raffini et al., 2025; Westram et al., 2021). Here, we study the flat periwinkle *Littorina fabalis*, which lives associated with seaweeds along the European Atlantic coast. This species has direct development with egg-masses attached to the seaweed, and a dispersal distance of about 8m per generation (Le Moan et al. 2024). It segregates into ecotypes associated with different levels of wave-exposure: In the NE Atlantic, a dwarf ecotype is located on shores sheltered from wave action, and a large ecotype inhabits shores with intermediate wave-exposure (Kemppainen et al., 2011; Reimchen, 1981; Tatarenkov and Johannesson, 1999). Although less studied, previous work also mentions additional ecotypes on the Atlantic coast of Iberia(Rolán and Templado, 1987), including a very small and reddish-coloured ecotype found among the red algae *Mastocarpus stellatus* in the most heavily wave-exposed shores. Demographic modeling suggests that the Swedish dwarf and large ecotypes have diverged in allopatry, but are now hybridizing in zones of secondary contact (Le Moan et al., 2024). Experimental manipulations and mating trials propose that size dimorphism is a result of local adaptation and weak assortative mating (Kemppainen et al., 2005; Saltin et al., 2013). A strong deficit in heterozygotes observed in Swedish hybrid zones furthermore suggests the presence of intrinsic genetic incompatibilities involved in reproductive isolation between the dwarf and large ecotypes (Tatarenkov and Johannesson, 1998). Recently, whole genome re-sequencing (WGS) of 295 *L. fabalis* sampled over 150 m parallel environmental gradients in Sweden showed that the genetic differences between the dwarf and large ecotypes clustered within 12 chromosomal inversions showing sharp arrangement frequency differences between moderately exposed and sheltered microenvironments (Le Moan et al., 2024, 2023). Both the nature of the barriers generated by the inversions, and their phenotypic consequences remain largely unknown.

This study compares two transects across similar environmental gradients of *Littorina fabalis*: the one in Sweden across the previously studied hybrid zone (Le Moan et al., 2023, 2024) and one in France. Previous work showed that these regions share substantial mitochondrial diversity, indicating common phylogeographic origins (Sotelo et al., 2020). In addition, population genetic studies based on one allozyme (arginine kinase) and on AFLP data revealed patterns compatible with parallel differentiation between sheltered and exposed habitats in France and Sweden (Galindo et al., 2021; Tatarenkov and Johannesson, 1999). Although we now know that the arginine kinase locus is located within a large chromosomal inversion on *L. fabalis* (Le Moan et al., 2023), direct evidence is lacking for the presence of these structural variants outside Sweden. Here, using whole-genome sequencing, we investigate the extent of phenotypic and genomic parallelism over similar wave-exposure gradients. These replicated transects allow us to assess whether chromosomal inversions harbour universal barrier loci that consistently restrict gene flow across contrasting shore environments, and/or whether they contribute to location-specific genetic differences. We further investigate the nature of any barriers to gene exchange associated with any inversions, starting with a test for reversed clines that would indicate intrinsic barrier loci within inversions. Then, using GWAS, we test whether these inversions influence size and colour, the two main phenotypic traits distinguishing the ecotypes (Kemppainen et al., 2005; Reimchen, 1981). Finding such associations would suggest that inversions protect locally adaptive alleles from recombination, maintaining ecotype-specific trait combinations under divergent selection.

## Materials and Methods

### Geographic and genomic sampling

We collected *Littorina fabalis* snails on brown and red algae along a Swedish and a French transect that spanned similar gradients of wave-exposure (Fig. 1). The Swedish samples (*n*=168 individuals) were collected in the spring of 2018 and their precise positions recorded by a Trimble total station along a 180 m transect on the island of Lökholmen, Sweden (58°53 N, 11°06 E, Fig. 1A). These samples, already analysed in Le Moan et al. (2024, 2023), were re-analysed here and compared with 137 individuals sampled using the same procedures during the spring and summer of 2022 along a 348 m transect in Lampaul-Plouarzel, on the west coast of France (48°46 N, 4°77 E, Fig. 1B). As much as possible, the aim was to sample the same ecological gradient from sheltered to exposed in each region.

**Figure 1:**
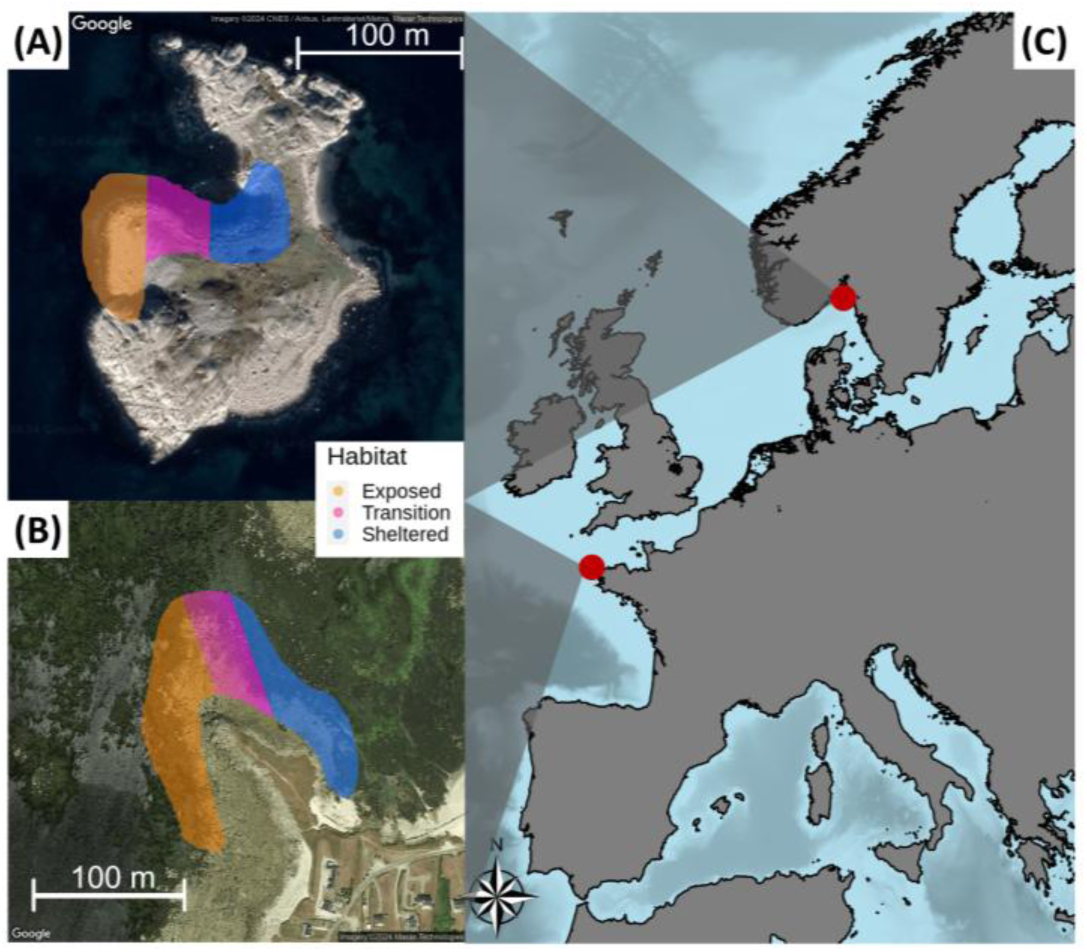
Location of sampling sites in Lökholmen, Sweden (A) and Lampaul-Plouarzel, France (B). The colour gradient represents the wave-exposure gradient, with three main zones (exposed, transition, sheltered) defined by the presence/absence of Ascophyllum nodosum, and in France also the presence of Mastocarpus stellus.

We characterized wave-exposure along our transects using algal communities as biological indicators (following Tatarenkov and Johannesson, 1994). The presence of *Ascophyllum nodosum* indicated sheltered areas, while its absence marked moderate to strong wave action. Sheltered and exposed zones were relatively distinct with a narrow transition zone in between (Fig. 1). In the French transect, *Mastocarpus stellatus* was present in the most exposed areas, but this species is absent in Sweden. We transformed the recorded two-dimensional position of each snail into the position in one dimension using a least-cost path approach Westram et al. (2021). Size and colour are the main phenotypic traits that vary with wave-exposure (Reimchen, 1981). We photographed each snail with its aperture facing upward alongside a scale bar. Using ImageJ software (Collins, 2007), we measured shell size as the maximum distance from aperture rim to apex. In addition, each individual was classified as yellow or brown and sexed. A piece of head/foot tissue from each individual was stored in 95% ethanol, from which we extracted DNA using the protocol of Panova et al., (2016). Individual Nextera whole genome sequencing libraries were prepared by SciLifeLab (Sweden) and sequenced on a NovaSeq S10000 with a minimum target coverage of 5X.

### Raw Data Bioinformatics Pipeline

The bioinformatics pipeline developed by Reeve et al. (2023) for short-reads WGS data from *Littorina* snails was used to process raw sequencing data. We automated and modified this protocol using a Snakemake (Mölder et al., 2025), available at https://github.com/PAJOT-Basile/Snakemake_processing_of_short-read_data. Specifically, raw sequencing reads were mapped to the new chromosome-level reference genome of the related species *Littorina saxatilis* (De Jode et al., 2024) using bwa-mem (Li and Durbin, 2009). Note that the reference genome used here is therefore an updated version of the one previously used in Le Moan et al. (2023, 2024). Mapped reads were then sorted, and duplicated reads were marked with samtools v1.15.1 (Li et al., 2009) before conducting single-nucleotide polymorphism (SNP) calling using bcftools mpileup (Li et al., 2009). Final filtering steps were performed using BCFtools (Danecek et al., 2021) and VCFTools (Danecek et al., 2011) to retain only biallelic SNPs located more than 5 bases away from an indel and exhibiting less than 5% of missing data and with a mean coverage between 5x and 13x. This dataset finally containing 16.1 million SNPs was further filtered depending on the specific requirements of each analysis, as described below.

### Phenotypic clines

The variation in shell size and shell colour over the transects, two traits that are involved in local adaption in *L. fabalis* (Kemppainen et al., 2005; Reimchen, 1981) were examined through cline analyses using the functions developed in Derryberry et al. (2014). We performed cline fitting using a maximum likelihood approach with the R package bbmle (Bolker, 2020). We used the quantitative trait cline function to investigate size variation, while we treated colour as dimorphic and fitted a cline to variation in the frequency of the brown morph as described in Westram et al. (2018). We tested the relevance of the clinal model by comparison with the expected results of a model with no variation and a linear variation model (Westram et al., 2018). As these models have different numbers of parameters, we selected the best model as the one with the lowest AIC. Significant differences between models were considered if the differences in AIC were greater than 4 (Westram et al., 2018).

### Population and ecotype genetic structure

We randomly selected one SNP every 1kb to reduce the effect of physical linkage of markers on population structure. In addition, to remove potential paralogs, we filtered out all SNPs with a heterozygosity rate greater than 0.55 (as performed in Le Moan et al., 2024). The genome-wide *F*_ST_ between French and Swedish populations was calculated with the 507k SNPs remaining after filtering using VCFTools (Danecek et al., 2011), following the method of Weir and Cockerham (1984). We used the same approach and SNP set to calculate the genome-wide *F*_ST_ values between ecotypes for each transect, but using 20 "reference" individuals from the sheltered end of each transect and 20 from the exposed end; these reference sets were used throughout subsequent analyses.

A principal component analysis (PCA) was then performed using the adegenet R package (Jombart, 2008). As principal components 2 and 3 both seemed to be associated with wave-exposure gradients in France and Sweden (see Results), we investigated these associations by plotting each individual’s PC2 and 3 coordinates against their position on the sampling transect. The individual contribution of each SNP to PC2 and 3 was then plotted along the genome as a first step to explore the distribution of genomic variation seemingly correlated to wave-exposure. Finally, PCA was run independently for the French and Swedish transects using a 500kb sliding window approach. This local PCA method allowed us to identify genomic regions associated with locally stronger than average genomic structure.

### Signatures of chromosomal rearrangements

The local PCA analyses revealed large chromosome regions displaying patterns consistent with chromosomal rearrangements. A previous study (Le Moan et al., 2024) identified these regions as associated with 12 putative inversions linked to ecotype differentiation, but that work focused exclusively on Swedish populations and relied on a fragmented *Littorina saxatilis* reference genome (Westram et al., 2018). Here, our study includes both Sweden and France, and we redefined these putative inversions using a new chromosome-level reference genome (De Jode et al., 2024). We (re-)identified and delineated inversion limits based on their indirect genomic signatures on differentiation, linkage disequilibrium (LD), and heterozygosity (Mérot et al., 2020; Reeve et al., 2024). To do so, we first recalculated SNP-wise *F*_ST_ between the reference ecotypes using only highly polymorphic SNPs from the raw dataset (16.1 million SNPs), retaining variants with a minor allele frequency ≥ 0.3 and without filtering for physical linkage. To identify genomic regions showing differentiation exceeding background levels, we applied a Hidden Markov Model (HMM) to the *F*_ST_ values for these highly polymorphic SNPs (Hofer et al., 2012; Marques et al., 2017; Soria-Carrasco et al., 2014). The HMM was run separately for each country. Within each country, a region was considered an inversion when it met four criteria: i) high differentiation according to the HMM; ii) a differentiated region at least 1 Mbp long; iii) three clear PCA clusters, with the central cluster positioned at mid-distance of the other two (Mérot et al., 2020); iv) higher heterozygosity in the central cluster than in the outer clusters. It is important to note that the filtering applied for the HMM excludes inversions with low minor-arrangement frequency, biasing detection toward inversions with strong ecotype differentiation, which aligns with the aims of our study. In criterion iii, the three PCA clusters correspond to the three possible karyotypes, with heterokaryotypes forming the central cluster. Clustering was performed using find.clust in adegenet (Jombart et al., 2010) with K = 3, and clusters were relabeled so that the most common cluster in exposed individuals was consistently designated as cluster 1. Heterozygosity for each cluster was calculated independently as the proportion of heterozygous SNPs within the inversion. Final confirmation of the inversions was performed by examining pairwise linkage disequilibrium (LD) along each transect using VCFTools (Danecek et al., 2011). LD was calculated independently using the same set of highly polymorphic SNPs (allele frequency ≥ 0.3) across all individuals, thinning to one SNP every 2.5 kb to reduce computational load. LD between all SNP pairs was visualized as heatmaps using ggplot2 (Wickham, 2016). To avoid artificial splitting of nearby inversions due to minor discrepancies between the *L. fabalis* genome and the *L. saxatilis* reference, three pairs of regions on the same chromosome that displayed identical PCA clustering patterns (LD=1) were merged (Fig. S11).

### The contribution of inversions to parallel and location-specific genetic differentiation between ecotypes

These analyses revealed that the genome is strongly structured by chromosomal inversions, which are strongly linked with ecotype differentiation. To investigate parallelism in inter-ecotype differentiation between Sweden and France, we first calculated the allelic frequency difference at each SNP between sheltered and exposed reference individuals as Δfreq = freq_sheltered – freq_exposed. We then computed the f4 statistic (Patterson et al., 2012) as f4 = Δfreq_France × Δfreq_Sweden. Positive f4 values indicate parallel differentiation, where the same alleles are consistently associated with either sheltered or exposed habitats in both countries. In contrast, negative f4 values indicate reverse differentiation, where SNPs show high Δfreq in both countries but with opposite allelic associations with the wave-exposure gradient (i.e., different alleles are associated with exposed habitats in each country). We then focused on the 5% most differentiated loci between ecotypes in each country to limit the computational time needed to assess the significance of Δfreq and f4 using permutation tests (n = 10,000 permutations). The permutations were performed by randomly shuffling the reference individuals from the sheltered and exposed habitats within each country independently. For each SNP, we tested the absolute allele frequency difference (|Δfreq|) in France and Sweden using one-tailed tests, and the f4 statistic using a two-tailed test to identify SNPs showing either private (|Δfreq|> 0 in one country but not in the other), parallel (|Δfreq|> 0 in both countries and f4 > 0) or reverse (|Δfreq|> 0 in both countries and f4 < 0) differentiation. P-values were calculated as the proportion of permutations yielding statistics as extreme as or more extreme than the observed values. To account for multiple testing across all SNPs, we applied FDR correction with significance thresholds of p < 0.05 for Δfreq and p < 0.025 for the f4 statistic.

To better understand the transect distribution of the putative inversions, we performed cline analyses using the genotypes of the inversions previously defined by local PCA (see Signatures of chromosomal rearrangements). Allele frequency cline functions that included direct estimates of FIS along the transects were fitted using RStan (Stan Development Team, 2024) as described in Le Moan et al. (2024). The FIS of the inversion genotypes assumed to peak at the cline center and estimated at 5 points, following the formulation in Cfit (Gay et al., 2008). The *F*_IS_ estimates in RStan made the models more suitable for genotype data than the classical maximum likelihood clines used earlier for colour. The degree of coincidence of the inversion clines, also known as coupling, was then measured using estimates of LD across inversion genotypes close to the zone centre (defined as the mean cline centre ± standard deviation of cline centres), and the coefficient of variation of the cline slopes, as suggested in (Firneno et al., 2023). Finally, genetic divergence between inversion arrangements and between countries was analyzed by constructing neighbor-joining (NJ) trees from homokaryotypic individuals. For each individual, one allele was randomly sampled from diploid genotypes to generate haploid sequences using a custom R script (available in the Zenodo archive). Genetic distances were calculated using the F81 model (Felsenstein, 1981) and used to build NJ trees with the R package phangorn (Schliep, 2011), which were visualized using ggtree (Yu et al., 2018). We generated one tree per inversion (16 trees in total) and included eight control regions outside inversions that represent the range of inversion sizes observed. Inversion boundaries were defined using the HMM approach and are detailed in Table S3 and control regions in Table S7.

### Genetic variation associated with shell phenotypes

We conducted a genome-wide association study (GWAS) to detect genomic regions associated with shell colour and size. These analyses were performed separately for the Swedish and French transects and corrected for both kinship and the structure between ecotypes using pairwise relatedness between individuals (Tan and Atkinson, 2023). Relatedness was calculated in VCFtools (Danecek et al., 2011) using the method by Yang et al. (2010)(see Fig. S1). The GWAS analyses were computed on all individuals using the filtered dataset of 507k SNPs using the statenGWAS R package (Bart-Jan and Kruijer, 2023).

## Results

### Phenotypic Clines

Snail size showed a clinal distribution along the wave-exposure gradient both in France and Sweden (Fig. 2, top panel), with clines of similar width in both countries (46.1 m in France, 51.1 m in Sweden; with width not significantly different from each other, Table S1). Average shell sizes differed significantly between sheltered and exposed sides in both transects (Table S1). Large ecotypes averaged 11.5 mm (95% CI: 11.2–11.9) in Sweden and 10.9 mm (95% CI: 10.6–11.2) in France, while dwarfs averaged 8.2 mm (95% CI: 7.1–8.9) and 8.4 mm (95% CI: 7.9–8.8), respectively. Remarkably, the cline was reversed in France compared to Sweden (Fig. 2A vs. B).

**Figure 2:**
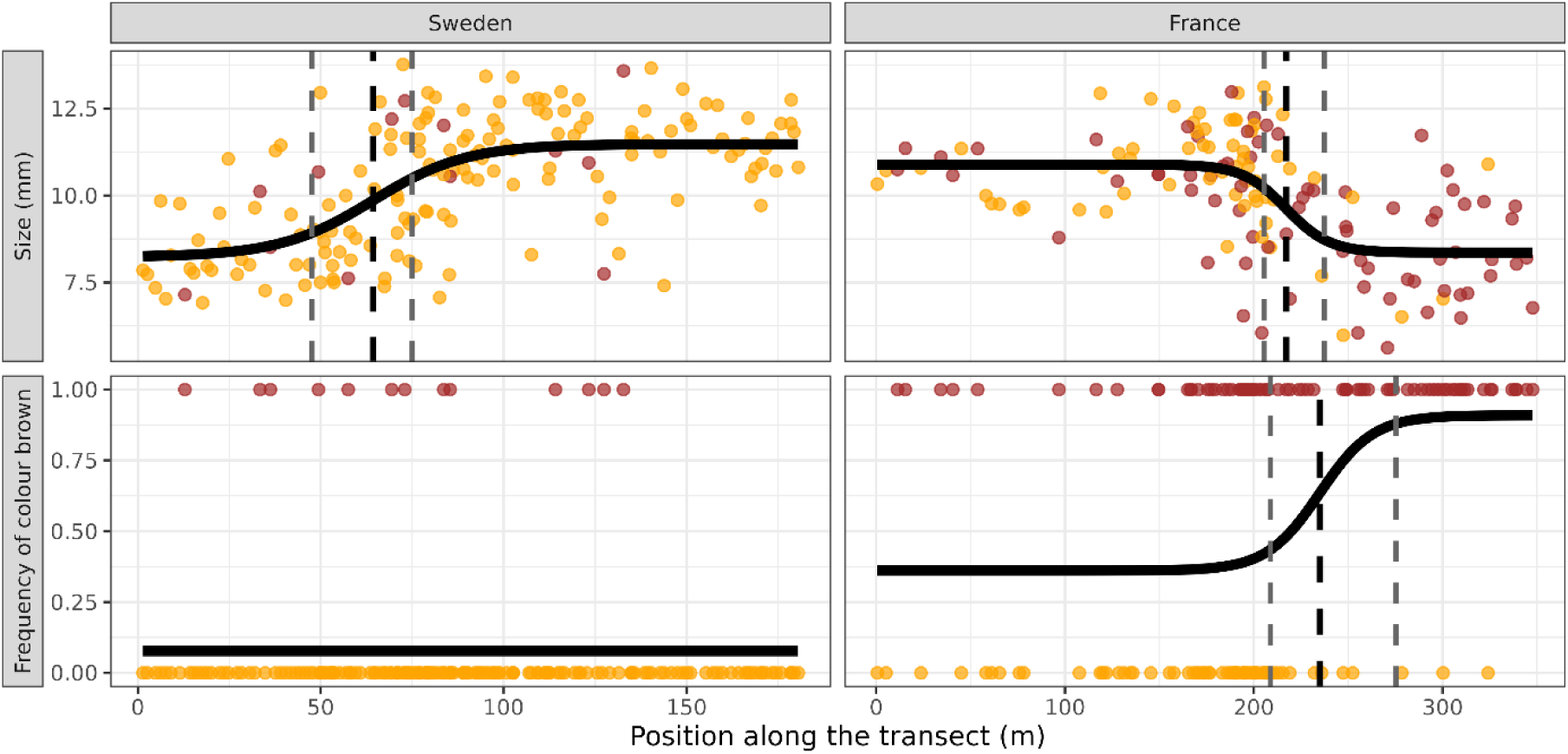
Distribution of Littorina fabalis shell phenotypes. along the Swedish and French transects. Both transects are sorted from the sheltered to the exposed part of the sea shore. Each point represents an individual, coloured according to its shell colour (yellow or brown). The top panels show the observed shell sizes at the Swedish (left) and French (right) sites and the bottom panels show the distribution of brown (y=1) and yellow individuals (y=0) along the Swedish (left) and French (right) transects. In all graphs, the black curve represents the best fitted phenotypic cline. The thick black dashed lines represent the centers of the phenotypic clines with their 95% confidence intervals represented by the dashed grey lines.

In addition, French snails showed greater variability in shell colour, with 45% yellow compared to 92% in Sweden (Fig. 2C-D). In Sweden, the few brown individuals (13) were spread along the transect (Fig. 2C), whereas in France brown individuals were mainly located in the exposed part of the transect. The center of the French colour cline (at 236 m) was slightly shifted towards the more exposed part of the transect compared to the size cline (216 m, Fig. 2, bottom panel), although this trend was not statistically significant (Table S1). For the colour variation in Sweden, it was not possible to distinguish the clinal model from the stable or the linear models (ΔAIC < 4), thus, the model with the fewest parameters (the stable model) was considered as the most likely model in Figure 2.

### Population structure

The PCA conducted to examine the spatial and ecological components of genetic structure in *L. fabalis* showed that the Swedish and French populations formed two distinct clusters (first axis, 10.59% variance explained, Fig. 3A). The mean *F*_ST_ between the Swedish and the French populations was 0.139 (p-value < 0.01). Axes 2 and 3, accounting for 5.16 % and 0.96 % of the total variation, respectively, were both correlated with wave-exposure (Figs. 3A-D). Individuals with intermediate PC values on these two axes were primarily located around the centers of the transects. These individuals also displayed strong heterozygosity, as expected for recent hybrids between the two ecotypes (Fig. S1). The mean *F*_ST_ value between ecotypes (i.e., using sheltered vs. exposed reference samples genotyped at 507k SNPs) was 0.16 in Sweden (p < 0.01) and 0.13 in France (p < 0.01). While both axes 2 and 3 were correlated with wave-exposure, the direction of this association was the same in Sweden and France for axis 2, but reversed for axis 3 (Figure 3 C,D). The third principal component thus captures genetic variation that seems correlated with the distribution of individuals along ecological transects, but with opposite orientations in France vs Sweden. High SNP loading scores on these axes were mostly clustering in some large genomic regions, and these regions appeared to be broadly the same for both axes (Fig. S3).

**Figure 3:**
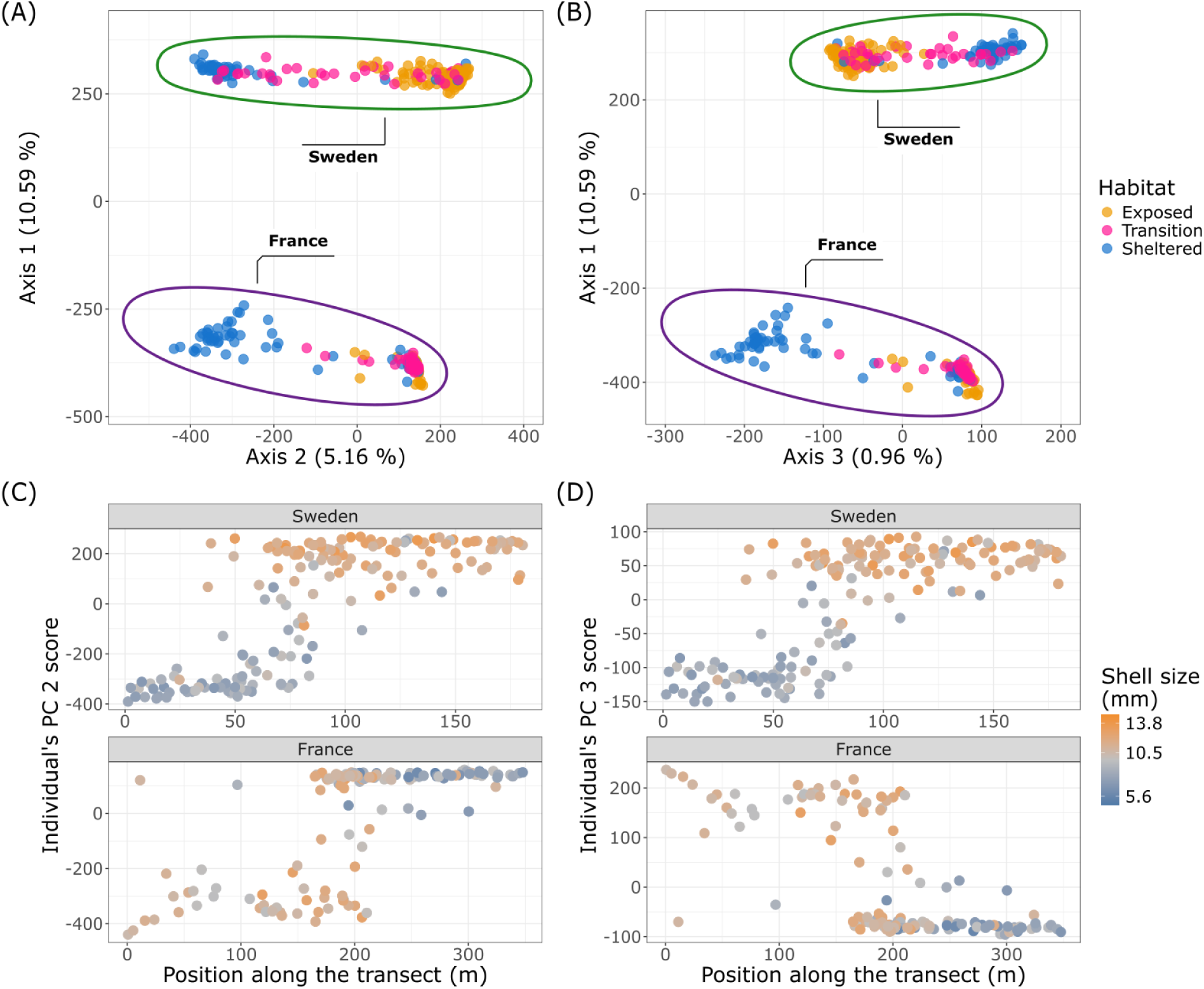
Population and ecotype structure of L. fabalis. (A) and (B) multilocus PCA based on 507k SNPs. Each dot is an individual, coloured according to its position along the wave-exposure gradient. (C) and (D) distribution of the individual coordinates on axes 2 and 3, respectively, relative to the individual positions along the transect. The colour of the points represents the size of the individuals from small (blue) to large (orange).

### Signatures of putative chromosomal inversions

The local PCA based on 500 kb sliding windows identified genomic regions associated with locally stronger than average genomic structure (Fig. 4A) and where samples were clearly split into three distinct groups. Importantly, these regions matched the putative inversions previously identified in Sweden (Le Moan et al. 2024), demonstrating that the same chromosomal rearrangements are present in both France and Sweden (Le Moan et al. 2024). Applying the four-criteria approach (Fig. S4-S8, Table S2) we identified 15 genomic regions in Sweden and 16 in France, in both cases including the 12 chromosomal inversions previously suggested by Le Moan et al. (2024). These inversions ranged from 1.5 Mbp to 56.5 Mbp (Table S3). With our new analysis using the high-quality reference genome (De Jode et al., 2024) we found two additional inversions not identified earlier, one on LG1 (shared by the two countries), one on LG2 private to the French population. Additionally, what were previously thought to be single inversions on LG6 and LG16 appeared to be two closely linked inversions in complete LD in Sweden but not in France (Fig. S9).

**Figure 4:**
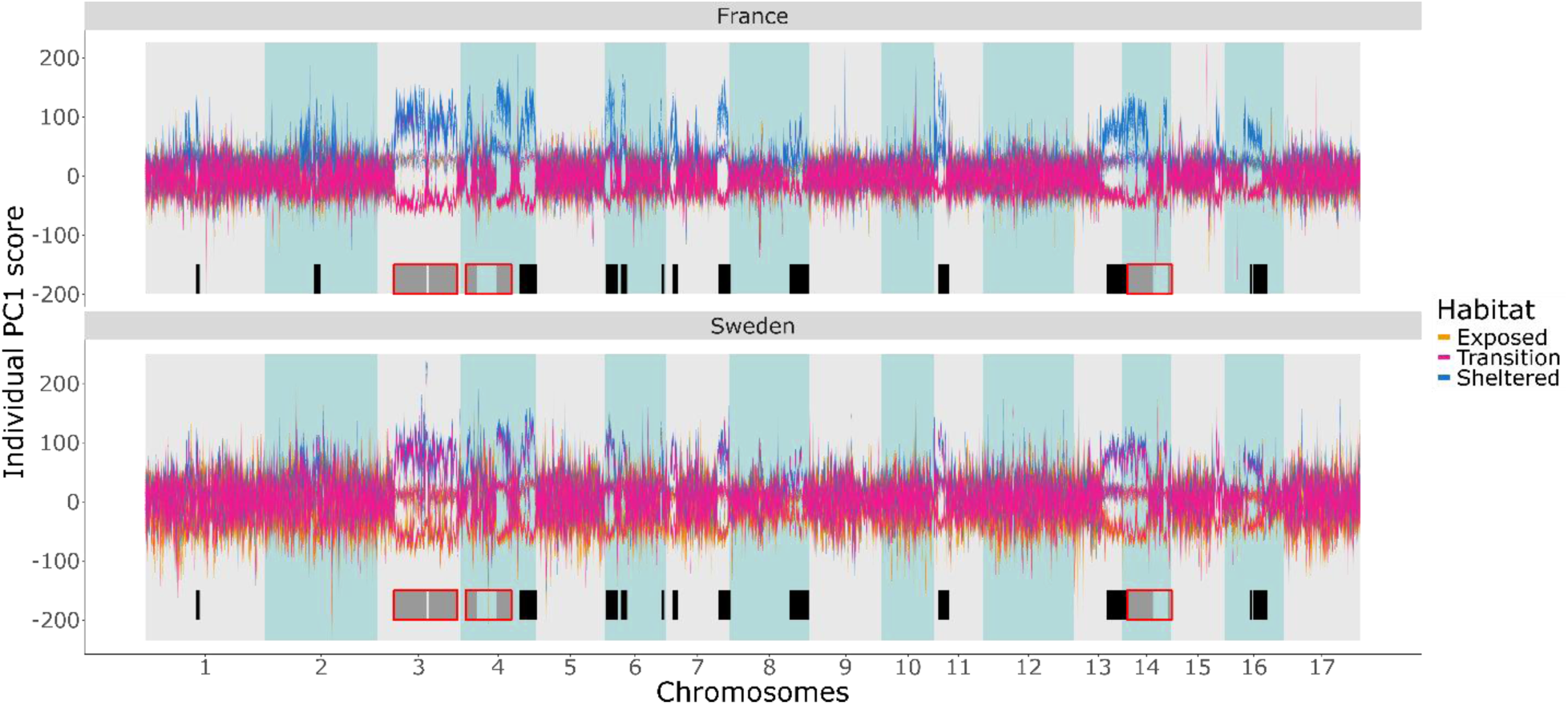
Local PCA analyses based on 500kb windows. using only French samples (panel A, top) and only Swedish samples (panel B, bottom), revealing the three sample clusters characteristic of chromosomal inversions. Each line represents one individual, coloured according to its habitat (sheltered in blue, transition in pink, and exposed in orange), and tracks the variation of individual coordinates on axis 1 of the sliding window PCA along the genome. Black boxes indicate inversion boundaries, and grey boxes with red outlines indicate inversions with complete linkage disequilibrium (LD = 1; Fig. S9).

### Genetic differentiation between ecotypes in France vs Sweden

The putative inversions harboured substantial genetic differences between ecotypes (with allele frequency differences between reference samples up to Δfreq = 1), which contributed to the marked heterogeneity in differentiation observed across the genome (Fig. 5). To identify the most differentiated regions and evaluate the extent of parallel and non-parallel ecotype differences between countries, we extracted SNPs in the top 5% quantile for allele frequency differences: 25,304 SNPs with Δfreq > 0.47 in Sweden and 24,500 SNPs with Δfreq > 0.45 in France. Combined, these represented 30,619 unique loci, 63% of which were shared between countries. Strikingly, 96% of these highly differentiated loci fell within the chromosomal inversions identified in Fig. 4. Among these differentiated loci, 88% (n = 26,917) exhibited parallel differentiation, with the same alleles contributing to ecotype differentiation in the same direction in both France and Sweden (both Δfreq and *f*4 statistics significantly > 0 after permutation test). The remaining outlier loci showed non-parallel patterns. These included country-specific outliers that showed significant differentiation in one country but no differentiation (Δfreq ≈ 0) in the other (1,720 private to France, 1,851 private to Sweden), as well as 131 loci with reversed inter-ecotype differentiation (both Δfreq and f4 significantly < 0 after permutation test). Like the parallel outliers, these non-parallel outlier loci were also concentrated within inversions, with 93% of reversed loci and 87–88% of private loci (in France and Sweden, respectively) located within the inversion regions. *F*_ST_ values between ecotypes reflected these patterns of differentiation. Parallel loci exhibited high and similar differentiation in both countries (Sweden mean *F*_ST_ = 0.76, France mean *F*_ST_ = 0.73), substantially exceeding values observed for reversed loci (Sweden mean *F*_ST_ = 0.40, France mean *F*_ST_ = 0.48). Country-specific outliers displayed strongly asymmetric *F*_ST_ patterns: French-private outliers showed high differentiation in France (*F*_ST_ = 0.58) but near-zero differentiation in Sweden (*F*_ST_ = 0.03), while Swedish-private outliers showed the reciprocal pattern (Sweden *F*_ST_ = 0.61, France *F*_ST_ = 0.03). Detailed pairwise *F*_ST_ matrices are provided in the supplementary table S5-S7.

**Figure 5:**
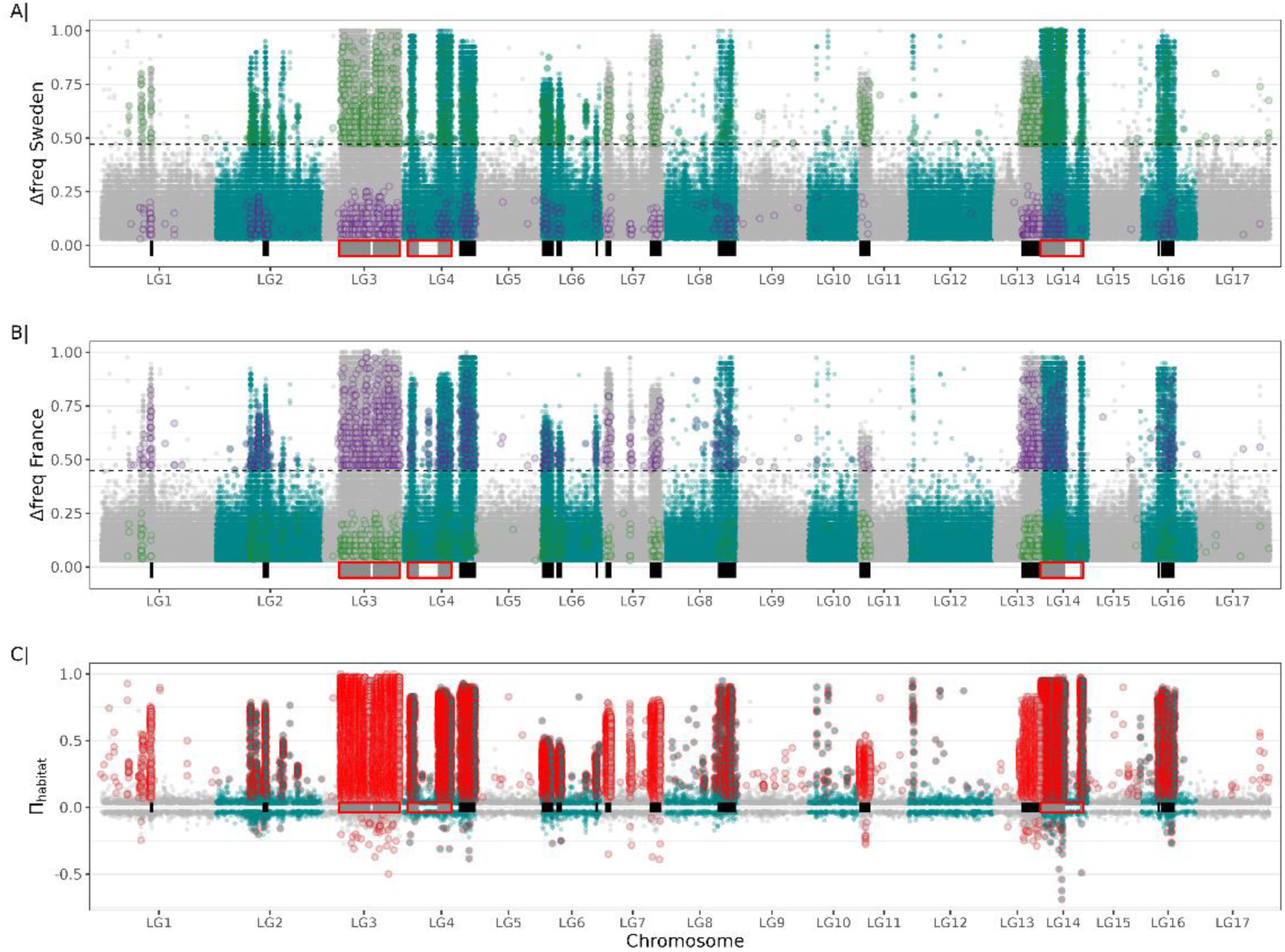
Ecotype differentiation in France and Sweden. (A-B) Distribution of absolute allele frequency differences (Δfreq) along the genome between exposed and sheltered reference samples in Sweden (A) and France (B). Dotted lines indicate the 95th percentile (0.47 and 0.45, respectively) used to identify highly-differentiated SNPs for permutation testing. Colored points represent private SNPs with significant habitat differentiation that are location-specific: purple for France-specific and green for Sweden-specific. (C) Genomic distribution of the Π*habitat* statistic. Positive values indicate parallel differentiation between exposed and sheltered ecotypes in both countries; negative values indicate reversed differentiation. SNP with a Πhabita significantly different from 0 after permutation testing are highlighted in red. Black boxes at the bottom of each plot denote inversion boundaries; red outlines indicate inversions in complete LD (LD = 1).

### The role of inversions in generating parallel and site-specific differences

The distribution of parallel outlier loci indicates that inversions drive most of the differentiation between ecotypes. Consistent with this pattern, cline analyses of inversion karyotypes—genotyped from local PCA—revealed striking similarities between the two countries. All inversions displayed steep clines with parallel frequency changes across countries, supporting the hypothesis that these inversions responded to selection along the wave-exposure gradients in a similar manner in both transects. All inversion cline centers were positioned at approximately the same location within each transect, averaging 63.25 m (SD = 10.48) in Sweden and 166.61 m (SD = 29.12) in France (Table S7). Cline widths were statistically not different between countries, with the exception of LG16, which exhibited a significantly steeper cline in Sweden than in France (Table S7). Most inversions exhibited significant heterozygote deficits (*F*_IS_ > 0), with only two exceptions (LG4.2 and LG6.1) showing *F*_IS_ values not significantly different from zero in either country (Table S7). LD between inversions at the zone centre (LD France = 0.39 ± 0.16, and LD Sweden =0.36 ± 0.14) and the coefficients of variation (CV) of cline slopes (CV = 0.24 in Sweden and CV = 0.29 in France) were both high in both countries, providing a proxy for strong coupling between inversions (Firneno et al., 2023; Le Moan et al., 2024). Taken together, the consistent cline characteristics across all inversions—including similar cline centers and slopes—combined with high LD between inversions at the hybrid zone centers, suggest that comparable levels of coupling contribute to maintaining parallel frequency clines in France and in Sweden.

Despite this broad parallelism, inversions also harbour substantial numbers of non-parallel outliers, thereby contributing to site-specific ecotype differences. This geographic differentiation within inversions is evident in PCA analyses. For instance, considering the inversions on LG6 and LG14, PC2 separates French from Swedish samples within one arrangement, while PC3 captures France-Sweden differentiation within the alternative arrangement (Figs. 7A, 7B). This pattern, where both inversion arrangements have accumulated site-specific genetic variation visible along different principal components, is consistent across all inversions (Fig. S10-S11), indicating that polymorphisms contributing to both ecotypic and geographic differences have accumulated within each arrangement. Linkage disequilibrium (LD) between SNP polymorphisms from within the inversions was stronger in Sweden (mean LD = 0.27) than in France (mean LD = 0.14) for 14 of the 15 inversions, further suggesting that site-specific processes have differentially shaped genetic diversity within inversions in the two countries. Phylogenetic analyses of homokaryotypic snails revealed clear clustering by arrangement, with homozygotes for the exposed-environment arrangement separating from those carrying the sheltered-habitat arrangement (Fig.7C, 7D). Geographic divergence between countries was also evident within each arrangement (Fig.7C, 7D). This within-arrangement divergence between countries was stronger than the geographic divergence observed in collinear regions (Fig. 7E, Table S6), as expected from the large proportion of private outliers found in the inverted regions. We also note that arrangement divergence was asymmetric between country: in 10 of 15 shared inversions, Swedish arrangements showed greater divergence from each other than their French counterparts (Table S6, e.g. for LG6 and 14 in Fig. 7C-D).

**Figure 6:**
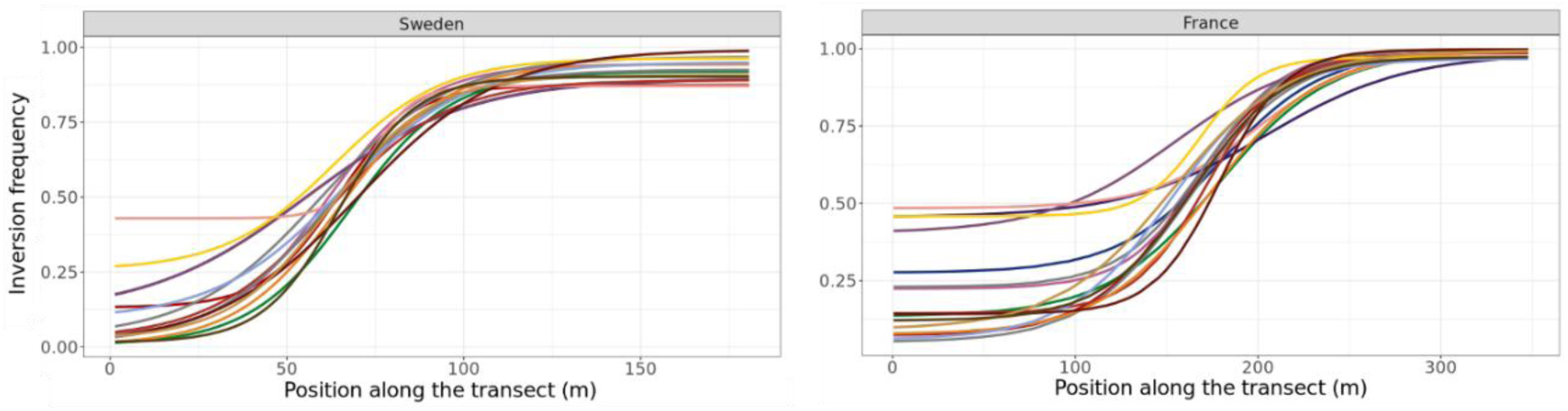
Frequency clines of the 16 chromosomal inversions for the Swedish transect (left) and the French transect (right). Each inversion cline is depicted in a different color (all inversion genotypes showed a significant clinal pattern).

**Figure 7:**
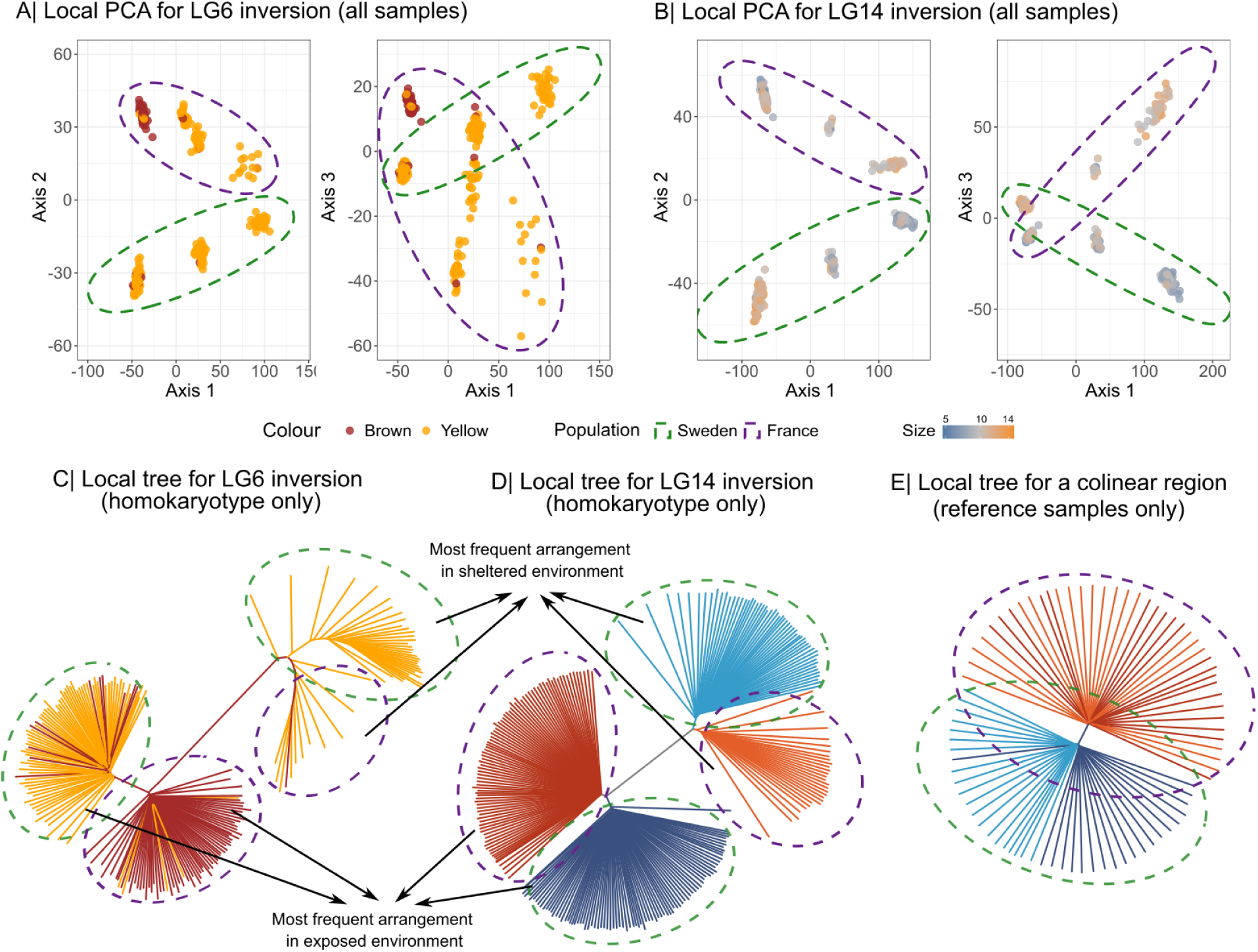
Detailed analyses of two chromosomal inversions (LG6.1 and LG14.1). A and B show the results of local PCA for Axis 1 vs. Axis 2 (left) and Axis 2 vs. Axis 3 (right). Axes 2 and 3 reveal substructures associated with alternative arrangements of the putative inversions. C and D show local phylogenetic trees for the two inversions, with a colinear comparison shown in E. For LG6, individuals are colored by shell color, as this inversion also carries the major color QTL (see below). The PCA for LG14 is colored by individual size (see color gradient, as this inversion also carries a size QTL), while the local tree is colored by sample karyotype. Local trees for colinear samples are colored by country (orange/red for France, blue for Sweden) and habitat (light for sheltered vs. dark for exposed). In each plot, French snails are encircled in purple, and Swedish snail in green.

### Genetic variation associated with shell phenotypes

In Sweden, GWAS identified a shell size QTL on LG14, with additional suggestive peaks on LG7 and LG16 (Fig. 8A). All three associations localized within inversions. For the LG14 QTL (inversion 14.1), local PCA confirmed the size association: PC1 (23.91% variance explained) separated smaller individuals (right cluster) from larger individuals (left cluster) within the Swedish samples (Fig. 7B, green ellipse). This pattern was Sweden-specific, as no genomic regions associated with shell size were detected in France. Inversion 14.1 also harboured SNPs showing reversed clines between locations (strong negative f4 values; Fig. 5A-B), with the five most extreme reversed SNPs located near the size-associated region (Fig. 5). These variants are good candidates for future work on the functional basis of the reversed association between shell size and wave-exposure observed across transects (Fig. 3).

**Figure 8:**
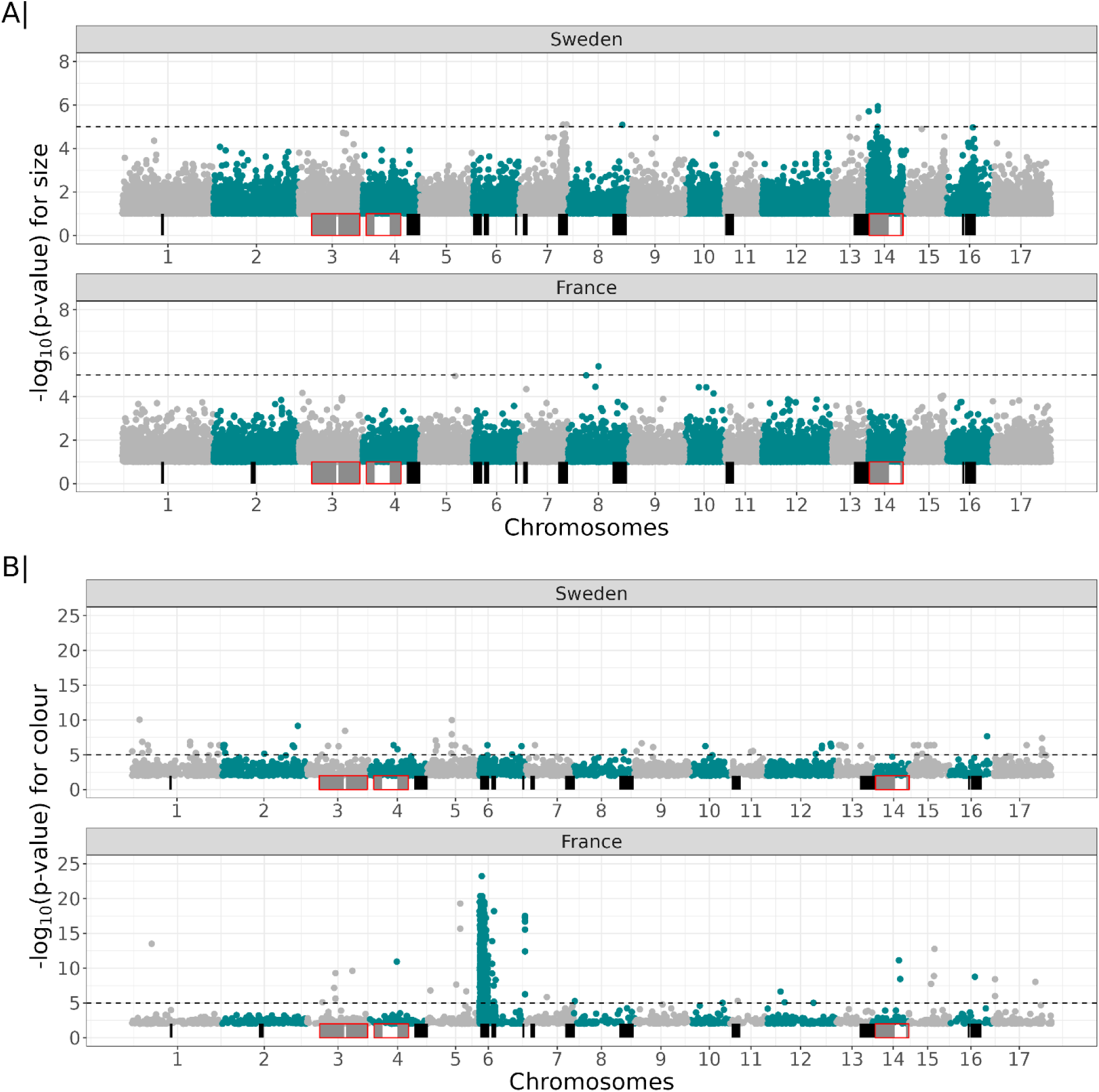
Genome-Wide Association Study (GWAS) on (A) shell size and on (B) shell colour. The Swedish and French populations were separated for the analysis. The dotted line represents the 10^-5^ p-value acceptance threshold value, and the black rectangles represent the identified chromosomal inversions.

GWAS on shell colour revealed a strong association with inversion 6.1 on LG6 in France (p < 10⁻²⁰; Fig. 8D). Polymorphisms within this inversion distinguished brown from yellow individuals in the local PCA (Fig. 7A, left cluster enriched for brown shells). Despite this inversion being polymorphic in Sweden, with 83.2% of the French colour-associated SNPs also variable there (Δfreq > 0.3), no significant colour QTLs were detected in the Swedish population (Fig. 8C, Fig. S7), consistent with limited colour variation at this site (Fig. 2A).

## Discussion

Genotypic differentiation between *L. saxatilis* ecotypes in wave-exposed and sheltered habitats appeared to be highly parallel across France and Sweden, with 96% of outlier loci concentrated within 15 shared inversions tightly coupled together in both countries. These inversions contain the bulk of the genetic basis underlying parallel ecotype differentiation in *Littorina fabalis*. Yet, despite this broadly parallel genomic architecture, our study reveals interesting discrepancies between genotypic and phenotypic patterns across the two hybrid zones. Colour variation shows a strong geographic contrast: brown morphs form a clear cline along the wave-exposure gradient in France but are nearly absent in Sweden, where 92% of individuals are yellow and homogeneously distributed across habitats (Fig. 2C-D). Shell size variation displays an even more surprising reversal pattern: large individuals occupy the more exposed habitats in Sweden but the more sheltered habitats in France (Fig. 2A-B). This genotype-phenotype decoupling is further exemplified by QTL mapping results. Notably, the association between inversion Inv_6.1 and colour was detected in France but absent in Sweden, despite this inversion being polymorphic and showing parallel clinal patterns at both sites. Similarly, the LG14 inversion is polymorphic in both countries, yet size QTLs are only detected in Sweden in this inversion, even though size is likely under selection in both countries. These findings demonstrate that the same inversions can harbour genetic variation that produces site-specific phenotypic responses. Understanding this phenomenon requires examining how local ecological condition, demographic history and selection on genetic variation within inversions jointly produce divergent phenotypes from a shared genomic architecture.

### The role of chromosomal inversions in the differentiation of ecotypes

The concentration of parallel genetic differentiation within chromosomal inversions aligns with growing evidence that structural variants play an important role in maintaining ecotype divergence over large geographic scales (Harringmeyer and Hoekstra, 2022; Jones et al., 2012; Matschiner et al., 2022; Meyer et al., 2024; Morales et al., 2019). Here, we detected 16 inversions using a high-quality reference genome (De Jode et al., 2024), including two relatively small inversions (1.6 and 2.3 Mbp) not identified in previous studies based on more fragmented assemblies (Le Moan et al., 2024, 2023). All shared inversions show large differentiation between ecotypes with parallel clines across transects. While such patterns of parallel differentiation are often interpreted as evidence of parallel adaptation from shared ancestral variation maintained by gene flow (Jones et al., 2012; Morales et al., 2019; Westram et al., 2022), previous demographic inferences on Swedish *L. fabalis* revealed that ecotypes evolved through allopatric divergence followed by secondary contact (Le Moan et al. 2024). The parallel genetic patterns we observe are therefore consistent with scenarios involving ecotype spread and secondary contact, though the exact sequence remains uncertain: ecotypes may have spread independently along European seashores before establishing contact zones in both France and Sweden, or alternatively, they may have spread from a single contact zone that later expanded. Either scenario could explain the similar levels of inversion coupling observed in both hybrid zones if epistatic interactions between inversions evolved during allopatric divergence, linking them together (as discussed in Le Moan et al., 2024) and thereby facilitating the parallel clines observed here. Similar patterns of replicated secondary contact zones have been documented in other marine coastal systems including European anchovies (Meyer et al., 2025), pipefish (Flanagan et al., 2021), isopods (Ribardière et al., 2025) and blue mussels (Fraïsse et al., 2018). Here, the ecotypic differentiation is highly heterogeneous along the genome in both countries: inversions harbour strongly differentiated regions with steep clines, while collinear regions show little differentiation. This pattern raises an important question about the evolutionary history of divergence and the timing of origin of the inversions: were collinear regions also strongly differentiated at secondary contact, with differences subsequently eroded by gene flow, or was divergence between allopatric ancestral populations already heterogeneously distributed and concentrated primarily within inversions? Distinguishing between these scenarios has important implications for understanding how inversions contribute to the origin versus the maintenance of ecotypic differentiation, but will require future work to be fully addressed.

While inversions carry most of the genetic polymorphism involved in parallel differentiation between ecotypes, they also concentrate more non-parallel loci (private and reversed outlier SNPs) than colinear regions. The weaker differentiation at these non-parallel SNPs (reversed SNP *F*_ST_ = 0.44; private SNP *F*_ST_ = 0.59) compared to parallel SNPs (*F*_ST_ = 0.74) suggests that they may reflect genetic drift acting more strongly within chromosomal inversions. Inversions typically have lower effective population sizes than collinear regions (Berdan et al., 2023), as inferred in *L. fabalis* (Le Moan et al., 2024), which intensifies the effects of drift. Site- and ecotype-specific demographic histories may also contribute to these patterns: the French and Swedish populations likely experienced different postglacial colonization dynamics, with the French site located closer to glacial refugia than the Swedish site, which only became available for marine life following deglaciation approximately 9,000 years ago (Björck, 1995). Genetic signatures from these distinct colonization histories (including associated founder events) are expected to persist longer in inversions where recombination is suppressed, while gene flow may erase them in collinear regions (Duranton et al., 2018). Altogether, these demographic processes could contribute to the site-specific genetic differences between ecotypes observed within inversions.

Importantly, not all site-specific differences between ecotypes can be attributed to demographic processes. In particular, the GWAS results show that at least 2 of the 15 shared inversions contain QTLs associated with shell colour and size that are specific to each locality. In the following sections, we discuss each of these phenotypic effects in detail, and how these site-specific phenotypic differences might relate to differing environmental conditions.

### Colour polymorphism and its variation across hybrid zones

Colour polymorphism in *L. fabalis* appears subject to geographically divergent selection pressures, as evidenced by a clear colour cline across the wave-exposure gradient in France but the absence of colour-environment associations in Sweden (Fig. 2). Although colour variation is not solely linked to local adaptation, having also been shown to contribute to disassortative mating in *L. fabalis* and thus to be subject to balancing selection (Gefaell et al., 2021), local ecological conditions can still play an important role in shaping morph frequencies. Previous studies have shown that different colour morphs provide camouflage against the predatory fish *Lipophrys pholis* in different algal habitats, with brown morphs favoured in exposed habitats and yellow morphs in sheltered habitats (Reimchen, 1979). Consistent with this mechanism, brown individuals in France were predominantly located in exposed habitats dominated by the dark-red alga *M. stellatus*, while yellow individuals were more common in sheltered areas with *F. vesiculosus* and *F. serratus* (two seaweed species that provide a background on which yellow *Littorina* shells may be less detectable to predators when located in the frond of the algae, Reimchen, 1979). In contrast, Sweden showed uniformly high frequencies of yellow morphs (92%) across the entire wave-exposure gradient, with the few brown individuals (n=13) distributed randomly along the transect. Although the very high frequency of yellow individuals in Sweden may partly reflect sampling bias, as yellow morphs are more conspicuous to human collectors than brown individuals, we are confident that the lack of a trend between environments in Sweden, but the association with the wave-exposure gradient in France, are not artifacts of this bias, as already an earlier study in Sweden did not find any colour by environment interaction (Ekendahl, 1994). The absence of such colour-environment associations in Sweden likely reflects differences in intertidal community composition between countries: both *M. stellatus* seaweeds and *L. pholis* fish are absent from Swedish sites, potentially relaxing selection for brown coloration. These geographic differences in community composition and associated colour variation demonstrate that the local ecological context has the potential to modulate selection, providing future opportunities to investigate the eco-evolutionary dynamics underlying the maintenance of polymorphism in hybrid zones.

The clear signature of the colour QTL detected in France is located in a large inversion at the beginning of LG6 (Fig. 6). Importantly, this genomic region is also covered by a colour-associated inversion in the closely related species *L. saxatilis* (Koch et al., 2021). Although colour variation differs between the two species (brown to yellow in *L. fabalis* vs. white-beige-black in *L. saxatilis*), these results suggest a shared genetic architecture for this trait at least between closely related *Littorina* species. In *L. fabalis* sampled in France, nearly all brown individuals (96%) were homokaryotypes for the most frequent in the exposed part of the shore, while heterokaryotypes or homokaryotypes for the alternative arrangement were yellow, with rare exceptions (possibly due to phenotyping errors). These results reveal important information about the genomic architecture of colour variation in the French population: the yellow allele(s) is likely located on the LG6 arrangement most common in sheltered habitats and is dominant over the brown allele(s), causing heterokaryotype individuals to be yellow. Conversely, the brown allele(s) is likely located on the arrangement most common in exposed habitats, and the brown colour is only expressed in homokaryotype individuals for the exposed arrangement. These kinds of simple genetic architectures, with clear dominance effects, appear to be relatively common in marine snails (Cole, 1975; Palmer, 1984; Gefaell et al, 2022).

Strikingly, this inversion is polymorphic and shows clinal frequency variation across the wave-exposure gradient in both France and Sweden, yet brown morphs were nearly absent from our Swedish samples, a pattern that demonstrates that inversions are not static entities with fixed allelic content. This has two important implications for understanding inversion dynamics in this system. First, because the arrangements at inversion 6.1 show genetic clines in both countries while colour shows a cline in France only, additional barrier traits under selection must be located on this inversion, maintaining clinal variation even where colour selection is relaxed. Second, the difference in colour allele frequencies between countries within the same arrangement demonstrates that inversion content can be shaped by local evolutionary processes. These findings challenge the view of inversions as static "supergenes" (Roesti et al., 2022) and instead support a more dynamic view where inversion arrangements serve as genomic vehicles upon which locally adaptive variation can accumulate independently across geographic regions (Faria et al., 2019).

### Why is variation in shell size across wave-exposure gradients reversed in Sweden and France?

Size is thought to be an adaptive trait responding to a combination of wave-exposure and predation (Reimchen, 1982). In Sweden, experimental translocation showed that larger sizes are better adapted to more wave exposed areas, due to the higher risk of dislodgement from the fucoid canopy: larger individuals have a better chance to survive predation when they fall from seaweeds into green crab territory (Kemppainen et al., 2005). In Sweden, snails in sheltered habitats reach a smaller size, similar to that observed in predator-free environments, as they can remain in the seaweed and are inaccessible to large crabs (Kemppainen et al., 2005). These associations between size and exposure/predation gradients have been described in populations from Sweden and Wales (Kemppainen et al., 2005; Reimchen, 1981). Yet the snail size cline in France is inverted compared to the clines in Wales and Sweden (pattern also observed in Spain, Galindo et al., 2021), suggesting that selection on shell size differs between locations. Green crabs were observed along the whole transect in Sweden (Kemppainen et al., 2005), but in France they are mostly present in the more sheltered habitat (A. Le Moan personal observations, and also observed elsewhere in the English Channel (Silva et al., 2010), which may explain why the size gradients are inverted between countries.

An alternative explanation is reversed coupling (Bierne et al., 2011), whereby stochastic processes during hybrid zone formation drive associations between endogenous barriers and environmental transitions. However, this hypothesis requires that shell size has no effect on fitness, or, acts as an endogenous barrier to gene flow (e.g., through assortative mating) rather than being under environmental selection—both assumptions seem unlikely (Saltin et al., 2013; Tatarenkov and Johannesson, 1998). Furthermore, our genetic analysis provided limited support for such strong, reversed differentiation patterns: we detected no reversion of inversion clines and only a very restricted number of SNPs (n = 134) showing strong, reversed differentiation between countries. This limited support for reversed coupling could be partially explained by the strong inversion coupling observed in both countries, suggesting that interactions between inversions constrain the reshuffling of chromosomal arrangements. Overall, these findings suggest that size variation more likely reflects differences in exogenous selection rather than endogenous processes.

This particular subset of SNPs showing strong inter-ecotype differentiation inverted between France and Sweden remains intriguing. Notably, five SNPs with very strong negative *f*4 value are located within the same inversion on LG14 that is also carrying the size QTL observed in Sweden (but not in France). These markers are possible candidate targets for selection that may contribute to the size variation gradient in both countries, including reversal of the gradient in one country. However, these GWAS results should be interpreted cautiously, as the size-associated SNPs were just above the significance threshold (unlike those for colour). The reason they were only detected in Sweden, despite size variation being present in both countries, is also unclear. Size is typically a highly polygenic trait, and this is true also in snails (Kess and Boulding, 2019; Koch et al., 2022), and for such traits, GWAS signals can be difficult to detect with low sample sizes (Hong and Park, 2012), as analyzed here. Additionally, genomic regions of low recombination, such as inversions, may also bias the detection of significant associations in GWAS analyses (Boyle and Noor, 2004). Nevertheless, previous QTL mapping studies in three natural populations of the sister species *L. saxatilis* also detected positive associations between size and an inversion on LG14 in two out of three studied hybrid zones (Koch et al., 2021), suggesting a shared genomic architecture of size in both systems.

## Conclusion

This study demonstrates that the pairs of *L. fabalis* ecotypes along Swedish and French shores share extensive genetic differentiation repeatedly associated with the wave-exposure gradient. These parallel differences between ecotypes are concentrated within 15 high-differentiation genomic regions displaying signatures characteristic of chromosomal inversions. However, despite this strong parallelism, the content of inversions shows important differences between the two hybrid zones. Recent theoretical work by Roesti et al., (2022) suggests that associations evolving within inversions might constrain local adaptation when initial selective conditions that favored particular associations of alleles differ between populations. However, our results suggest that fine-tuned adaptative processes may occur. By demonstrating substantial variation in QTL content between two spatially separated hybrid zones that nevertheless show important genetic parallelism, we found that associations between loci within inversions can evolve differently between populations. These findings suggest that inversions are not entirely static entities but rather dynamic genomic scaffolds that can accumulate site-specific variation in response to local selection pressures (Berdan et al., 2023; Faria et al., 2019). Despite the strong barrier effects detected for most inversions, the vast majority of the traits under selection, and the mechanisms contributing to reproductive isolation between ecotypes, within these inverted regions remain largely unknown. Future experimental work is essential to understand how these differences evolved, and how contrasting demographic histories and variation in ecological communities across sites have shaped the evolution of inversion content and the traits they harbour.

## Acknowledgement

We are grateful to the many people who contributed to this work, particularly James Reeves and Maël Gross for their help with sampling, Olga Ortega-Martinez for DNA extractions, Charlotte Berthelier for her guidance on Snakemake, Jacques David for advice on GWAS analyses, and Sean Stankowski, Nicolas Bierne, the *Littorina* research community, Cécile Molinier, Claire Mérot, Pierre-Alexandre Gagnaire, and the DiSEEM team for valuable discussions throughout the project. This work was supported by access to the EMBRC-France and Biogenouest Genomer platform at the Station Biologique de Roscoff and the Roscoff Bioinformatics platform ABiMS. We also thank the Station Biologique de Roscoff for providing internal funding to support Basile Pajot during his internship.

## Data availability statement

Raw fasta file for the WGS sequencing are available with NCBI SRA under the project name “*Littorina fabalis* transect WGS” with accession number PRJNA836378 for the swedish transect, and with this accession number “XXXX” for the french transect. The bioinformatic pipeline packaged into a snakemake is available here: https://github.com/PAJOT-Basile/Snakemake_processing_of_short-read_data. Other scripts use to produce the different results of the manuscript are available as a zenodo archive with the following DOI:

## Authors contributions

Conceptualization: ALM with advice from T.B., R.B., K.J, and R.F., sampling: A.L.M., R.F., T.B. and K.J., phenotyping: A.L.M. and K.J., bioinformatic and data analyses: B.P. and ALM, supervision: A.L.M. and T.B., Writing—original draft: B.P., A.L.M. and T.B., Writing—review and editing: all co-authors

## References

1. Berdan, E.L., Barton, N.H., Butlin, R., Charlesworth, B., Faria, R., Fragata, I., Gilbert, K.J., Jay, P., Kapun, M., Lotterhos, K.E., Mérot, C., Durmaz Mitchell, E., Pascual, M., Peichel, C.L., Rafajlović, M., Westram, A.M., Schaeffer, S.W., Johannesson, K., Flatt, T., 2023. How chromosomal inversions reorient the evolutionary process. J. Evol. Biol. 36, 1761–1782. 10.1111/jeb.14242

2. Bierne, N., Gagnaire, P.-A., David, P., 2013. The geography of introgression in a patchy environment and the thorn in the side of ecological speciation. Curr. Zool. 59, 72–86. 10.1093/czoolo/59.1.72

3. Bierne, N., Welch, J., Loire, E., Bonhomme, F., David, P., 2011. The coupling hypothesis: why genome scans may fail to map local adaptation genes. Mol. Ecol. 20, 2044–2072. 10.1111/j.1365-294X.2011.05080.x

4. Björck, S., 1995. A review of the history of the Baltic Sea, 13.0-8.0 ka BP. Quat. Int. - QUATERN INT 27, 19–40. 10.1016/1040-6182(94)00057-C

5. Boyle, A.S., Noor, M.A.F., 2004. Variation in recombination rate may bias human genetic disease mapping studies. Genetica 122, 245–252. 10.1007/s10709-004-1703-6

6. Butlin, R.K., Smadja, C.M., 2018. Coupling, Reinforcement, and Speciation. Am. Nat. 191, 155–172. 10.1086/695136

7. Cole T. J., 1975. Inheritance of juvenile shell colour of the oyster drill Urosalpinx cinerea. Nature, 257(5529), 794–795. 10.1038/257794a0

8. Collins, T.J., 2007. ImageJ for microscopy. BioTechniques 43, 25–30. 10.2144/000112517

9. Danecek, P., Auton, A., Abecasis, G., Albers, C.A., Banks, E., DePristo, M.A., Handsaker, R.E., Lunter, G., Marth, G.T., Sherry, S.T., McVean, G., Durbin, R., 1000 Genomes Project Analysis Group, 2011. The variant call format and VCFtools. Bioinformatics 27, 2156–2158. 10.1093/bioinformatics/btr330

10. Danecek, P., Bonfield, J.K., Liddle, J., Marshall, J., Ohan, V., Pollard, M.O., Whitwham, A., Keane, T., McCarthy, S.A., Davies, R.M., Li, H., 2021. Twelve years of SAMtools and BCFtools. GigaScience 10, giab008. 10.1093/gigascience/giab008

11. De Jode, A., Faria, R., Formenti, G., Sims, Y., Smith, T.P., Tracey, A., Wood, J.M.D., Zagrodzka, Z.B., Johannesson, K., Butlin, R.K., Leder, E.H., 2024. Chromosome-scale Genome Assembly of the Rough Periwinkle Littorina saxatilis. Genome Biol. Evol. 16, evae076. 10.1093/gbe/evae076

12. Derryberry, E.P., Derryberry, G.E., Maley, J.M., Brumfield, R.T., 2014. hzar: hybrid zone analysis using an R software package. Mol. Ecol. Resour. 14, 652–663. 10.1111/1755-0998.12209

13. Duranton, M., Allal, F., Fraïsse, C., Bierne, N., Bonhomme, F., Gagnaire, P.-A., 2018. The origin and remolding of genomic islands of differentiation in the European sea bass. Nat. Commun. 9, 2518. 10.1038/s41467-018-04963-6

14. Faria, R., Johannesson, K., Butlin, R.K., Westram, A.M., 2019. Evolving Inversions. Trends Ecol. Evol. 34, 239–248. 10.1016/j.tree.2018.12.005

15. Faria, R., Navarro, A., 2010. Chromosomal speciation revisited: rearranging theory with pieces of evidence. Trends Ecol. Evol. 25, 660–669. 10.1016/j.tree.2010.07.008

16. Firneno, T.J., Semenov, G., Dopman, E.B., Taylor, S.A., Larson, E.L., Gompert, Z., 2023. Quantitative Analyses of Coupling in Hybrid Zones. Cold Spring Harb. Perspect. Biol. a041434. 10.1101/cshperspect.a041434

17. Flanagan, S.P., Rose, E., Jones, A.G., 2021. The population genomics of repeated freshwater colonizations by Gulf pipefish. Mol. Ecol. 30, 1672–1687. 10.1111/mec.15841

18. Fraïsse, C., Roux, C., Gagnaire, P.-A., Romiguier, J., Faivre, N., Welch, J.J., Bierne, N., 2018. The divergence history of European blue mussel species reconstructed from Approximate Bayesian Computation: the effects of sequencing techniques and sampling strategies. PeerJ 6, e5198. 10.7717/peerj.5198

19. Galindo, J., Carvalho, J., Sotelo, G., Duvetorp, M., Costa, D., Kemppainen, P., Panova, M., Kaliontzopoulou, A., Johannesson, K., Faria, R., 2021. Genetic and morphological divergence between Littorina fabalis ecotypes in Northern Europe. J. Evol. Biol. 34, 97–113. 10.1111/jeb.13705

20. Gay, L., Crochet, P.-A., Bell, D.A., Lenormand, T., 2008. Comparing Clines on Molecular and Phenotypic Traits in Hybrid Zones: A Window on Tension Zone Models. Evolution 62, 2789–2806. 10.1111/j.1558-5646.2008.00491.x

21. Hager, E.R., Harringmeyer, O.S., Wooldridge, T.B., Theingi, S., Gable, J.T., McFadden, S., Neugeboren, B., Turner, K.M., Jensen, J.D., Hoekstra, H.E., 2022. A chromosomal inversion contributes to divergence in multiple traits between deer mouse ecotypes. Science 377, 399–405. 10.1126/science.abg0718

22. Harringmeyer, O.S., Hoekstra, H.E., 2022. Chromosomal inversion polymorphisms shape the genomic landscape of deer mice. Nat. Ecol. Evol. 6, 1965–1979. 10.1038/s41559-022-01890-0

23. Ekendahl, A., 1994. Factors important to the distribution of colour morphs of *Littorina Mariae* Sacchi & Rastelli in a non-tidal area. Ophelia 40:1–12

24. Gefaell, J., Galindo, J., Malvido, C. et al. (2021). Negative assortative mating and maintenance of shell colour polymorphism in *Littorina* (*Neritrema*) species. Mar Biol 168, 151. 10.1007/s00227-021-03959-z

25. Gefaell, J., Galindo, J., & Rolán-Alvarez, E., 2022. Shell color polymorphism in marine gastropods. Evolutionary applications, 16(2), 202–222. 10.1111/eva.13416

26. Hofer, T., Foll, M., Excoffier, L., 2012. Evolutionary forces shaping genomic islands of population differentiation in humans. BMC Genomics 13, 107. 10.1186/1471-2164-13-107

27. Hong, E.P., Park, J.W., 2012. Sample Size and Statistical Power Calculation in Genetic Association Studies. Genomics Inform. 10, 117–122. 10.5808/GI.2012.10.2.117

28. Huang, K., Andrew, R.L., Owens, G.L., Ostevik, K.L., Rieseberg, L.H., 2020. Multiple chromosomal inversions contribute to adaptive divergence of a dune sunflower ecotype. Mol. Ecol. 29, 2535–2549. 10.1111/mec.15428

29. Johannesson, K., 2003. Evolution in Littorina: ecology matters. J. Sea Res., Structuring Factors of Shallow Marine Coastal Communities, Part II 49, 107–117. 10.1016/S1385-1101(02)00218-6

30. Johannesson, K., Faria, R., Moan, A.L., Rafajlović, M., Westram, A.M., Butlin, R.K., Stankowski, S., 2024. Diverse pathways to speciation revealed by marine snails. Trends Genet. 40, 337–351. 10.1016/j.tig.2024.01.002

31. Johannesson, K., Panova, M., Kemppainen, P., André, C., Rolán-Alvarez, E., Butlin, R.K., 2010. Repeated evolution of reproductive isolation in a marine snail: unveiling mechanisms of speciation. Philos. Trans. R. Soc. B Biol. Sci. 365, 1735–1747. 10.1098/rstb.2009.0256

32. Jombart, T., 2008. adegenet: a R package for the multivariate analysis of genetic markers. Bioinformatics 24, 1403–1405. 10.1093/bioinformatics/btn129

33. Jombart, T., Devillard, S., Balloux, F., 2010. Discriminant analysis of principal components: a new method for the analysis of genetically structured populations. BMC Genet. 11, 94. 10.1186/1471-2156-11-94

34. Jones, F.C., Grabherr, M.G., Chan, Y.F., Russell, P., Mauceli, E., Johnson, J., Swofford, R., Pirun, M., Zody, M.C., White, S., Birney, E., Searle, S., Schmutz, J., Grimwood, J., Dickson, M.C., Myers, R.M., Miller, C.T., Summers, B.R., Knecht, A.K., Brady, S.D., Zhang, H., Pollen, A.A., Howes, T., Amemiya, C., Lander, E.S., Di Palma, F., Lindblad-Toh, K., Kingsley, D.M., 2012. The genomic basis of adaptive evolution in threespine sticklebacks. Nature 484, 55–61. 10.1038/nature10944

35. Kemppainen, P., Lindskog, T., Butlin, R., Johannesson, K., 2011. Intron sequences of arginine kinase in an intertidal snail suggest an ecotype-specific selective sweep and a gene duplication. Heredity 106, 808–816. 10.1038/hdy.2010.123

36. Kemppainen, P., Nes, S. van Ceder, C., Johannesson, K., 2005. Refuge function of marine algae complicates selection in an intertidal snail. Oecologia 143, 402–411. 10.1007/s00442-004-1819-5

37. Kess, T., Boulding, E.G., 2019. Genome-wide association analyses reveal polygenic genomic architecture underlying divergent shell morphology in Spanish Littorina saxatilis ecotypes. Ecol. Evol. 9, 9427–9441. 10.1002/ece3.5378

38. Koch, E.L., Morales, H.E., Larsson, J., Westram, A.M., Faria, R., Lemmon, A.R., Lemmon, E.M., Johannesson, K., Butlin, R.K., 2021. Genetic variation for adaptive traits is associated with polymorphic inversions in Littorina saxatilis. Evol. Lett. 5, 196–213. 10.1002/evl3.227

39. Koch, E.L., Ravinet, M., Westram, A.M., Johannesson, K., Butlin, R.K., 2022. Genetic architecture of repeated phenotypic divergence in Littorina saxatilis ecotype evolution. Evolution 76, 2332–2346. 10.1111/evo.14602

40. Le Moan, A., Gagnaire, P.-A., Bonhomme, F., 2016. Parallel genetic divergence among coastal–marine ecotype pairs of European anchovy explained by differential introgression after secondary contact. Mol. Ecol. 25, 3187–3202. 10.1111/mec.13627

41. Le Moan, A., Panova, M., De Jode, A., Ortega-Martinez, O., Duvetorp, M., Faria, R., Butlin, R., Johannesson, K., 2023. An allozyme polymorphism is associated with a large chromosomal inversion in the marine snail Littorina fabalis. Evol. Appl. 16, 279–292. 10.1111/eva.13427

42. Le Moan, A., Stankowski, S., Rafajlović, M., Ortega-Martinez, O., Faria, R., Butlin, R.K., Johannesson, K., 2024. Coupling of twelve putative chromosomal inversions maintains a strong barrier to gene flow between snail ecotypes. Evol. Lett. 8, 575–586. 10.1093/evlett/qrae014

43. Lenormand, T., 2002. Gene flow and the limits to natural selection. Trends Ecol. Evol. 17, 183–189. 10.1016/S0169-5347(02)02497-7

44. Li, H., Durbin, R., 2009. Fast and accurate short read alignment with Burrows-Wheeler transform. Bioinforma. Oxf. Engl. 25, 1754–1760. 10.1093/bioinformatics/btp324

45. Li, H., Handsaker, B., Wysoker, A., Fennell, T., Ruan, J., Homer, N., Marth, G., Abecasis, G., Durbin, R., 2009. The Sequence Alignment/Map format and SAMtools. Bioinformatics 25, 2078–2079. 10.1093/bioinformatics/btp352

46. Marques, D.A., Lucek, K., Haesler, M.P., Feller, A.F., Meier, J.I., Wagner, C.E., Excoffier, L., Seehausen, O., 2017. Genomic landscape of early ecological speciation initiated by selection on nuptial colour. Mol. Ecol. 26, 7–24. 10.1111/mec.13774

47. Matschiner, M., Barth, J.M.I., Tørresen, O.K., Star, B., Baalsrud, H.T., Brieuc, M.S.O., Pampoulie, C., Bradbury, I., Jakobsen, K.S., Jentoft, S., 2022. Supergene origin and maintenance in Atlantic cod. Nat. Ecol. Evol. 6, 469–481. 10.1038/s41559-022-01661-x

48. Mérot, C., Oomen, R.A., Tigano, A., Wellenreuther, M., 2020. A Roadmap for Understanding the Evolutionary Significance of Structural Genomic Variation. Trends Ecol. Evol. 35, 561–572. 10.1016/j.tree.2020.03.002

49. Meyer, L., Barry, P., Le Moan, A., Arbiol, C., Castilho, R., Van der Lingen, C., Chlaïda, M., McKeown, N., Ernande, B., Bonhomme, F., Gagnaire, P.-A., Guinand, B., 2025. Genome divergence between European anchovy ecotypes fuelled by structural variants originating from trans-equatorial admixture. Proc. R. Soc. B Biol. Sci. 292, 20251416. 10.1098/rspb.2025.1416

50. Meyer, L., Barry, P., Riquet, F., Foote, A., Der Sarkissian, C., Cunha, R.L., Arbiol, C., Cerqueira, F., Desmarais, E., Bordes, A., Bierne, N., Guinand, B., Gagnaire, P.-A., 2024. Divergence and gene flow history at two large chromosomal inversions underlying ecotype differentiation in the long-snouted seahorse. Mol. Ecol. 33, e17277. 10.1111/mec.17277

51. Mölder, F., Jablonski, K.P., Letcher, B., Hall, M.B., Dyken, P.C. van, Tomkins-Tinch, C.H., Sochat, V., Forster, J., Vieira, F.G., Meesters, C., Lee, S., Twardziok, S.O., Kanitz, A., VanCampen, J., Malladi, V., Wilm, A., Holtgrewe, M., Rahmann, S., Nahnsen, S., Köster, J., 2025. Sustainable data analysis with Snakemake. 10.12688/f1000research.29032.3

52. Morales, H.E., Faria, R., Johannesson, K., Larsson, T., Panova, M., Westram, A.M., Butlin, R.K., 2019. Genomic architecture of parallel ecological divergence: Beyond a single environmental contrast. Sci. Adv. 5, eaav9963. 10.1126/sciadv.aav9963

53. Nicolas, L.A., Berdan, E.L., Wellenreuther, M., Colinet, H., Clouard, A., Wit, P.D., Glémin, S., Mérot, C., 2025. Parallel clines of chromosomal inversion frequencies in seaweed flies are associated with thermal variation. 10.1101/2025.04.28.650981

54. Pal, A., Shipilina, D., Le Moan, A., McNairn, A.J., Grenier, J.K., Kucka, M., Coop, G., Chan, Y.F., Barton, N.H., Field, D.L., Stankowski, S., 2025. Genealogical Analysis of Replicate Flower Colour Hybrid Zones in Antirrhinum. Mol. Ecol. 34, e70067. 10.1111/mec.70067

55. Palmer, R. A., 1984. Species cohesiveness and genetic control of shell colour and form in Thais emarginata (Prosobranchia, Muricacea): Preliminary results. Malacologia, 25, 477–491.

56. Panova, M., Aronsson, H., Cameron, R.A., Dahl, P., Godhe, A., Lind, U., Ortega-Martinez, O., Pereyra, R., Tesson, S.V.M., Wrange, A.-L., Blomberg, A., Johannesson, K., 2016. DNA Extraction Protocols for Whole-Genome Sequencing in Marine Organisms. Methods Mol. Biol. Clifton NJ 1452, 13–44. 10.1007/978-1-4939-3774-5_2

57. Patterson, N., Moorjani, P., Luo, Y., Mallick, S., Rohland, N., Zhan, Y., Genschoreck, T., Webster, T., Reich, D., 2012. Ancient admixture in human history. Genetics 192, 1065–1093. 10.1534/genetics.112.145037

58. Raffini, F., De Jode, A., Johannesson, K., Faria, R., Zagrodzka, Z.B., Westram, A.M., Galindo, J., Rolán-Alvarez, E., Butlin, R.K., 2025. Phenotypic Divergence and Genomic Architecture Between Parallel Ecotypes at Two Different Points on the Speciation Continuum in a Marine Snail. Mol. Ecol. 34, e70025. 10.1111/mec.70025

59. Ravinet, M., Faria, R., Butlin, R.K., Galindo, J., Bierne, N., Rafajlović, M., Noor, M. A. F., Mehlig, B., Westram, A.M., 2017. Interpreting the genomic landscape of speciation: a road map for finding barriers to gene flow. J. Evol. Biol. 30, 1450–1477. 10.1111/jeb.13047

60. Reeve, J., Butlin, R.K., Koch, E.L., Stankowski, S., Faria, R., 2024. Chromosomal inversion polymorphisms are widespread across the species ranges of rough periwinkles (Littorina saxatilis and L. arcana). Mol. Ecol. 33, e17160. 10.1111/mec.17160

61. Reimchen, T., 1981. Microgeographical variation in Littorina mariae Sacchi & Rastelli and a taxonomic consideration. J. Conchol. 30, 341–350.

62. Reimchen, T.E., 1982. Shell size divergence in Littorina mariae and L. obtusata and predation by crabs. Can. J. Zool. 60, 687–695. 10.1139/z82-098

63. Reimchen, T.E., 1979. Substratum heterogeneity, crypsis, and colour polymorphism in an intertidal snail (Littorina mariae). Can. J. Zool. 57, 1070–1085. 10.1139/z79-135

64. Ribardière, A., Daguin-Thiébaut, C., Coudret, J., Corguillé, G.L., Avia, K., Houbin, C., Loisel, S., Gagnaire, P.-A., Broquet, T., 2025. Sex chromosomes and chromosomal rearrangements are key to behavioural sexual isolation in Jaera albifrons marine isopods. 10.1101/2025.01.08.631900

65. Riginos, C., Cunningham, C.W., 2005. Local adaptation and species segregation in two mussel (Mytilus edulis x Mytilus trossulus) hybrid zones. Mol. Ecol. 14, 381–400. 10.1111/j.1365-294X.2004.02379.x

66. Roesti, M., Gilbert, K.J., Samuk, K., 2022. Chromosomal inversions can limit adaptation to new environments. Mol. Ecol. 31, 4435–4439. 10.1111/mec.16609

67. Rolán Mosquera, E., Templado González, J., 1987. Consideraciones sobre el complejo Littorina obtusata-mariae (Mollusca, Gastropoda, Littorinidae) en el Noroeste de la Península Ibérica. Thalass. Int. J. Mar. Sci. 5, 71–85.

68. Rougemont, Q., Gagnaire, P.-A., Perrier, C., Genthon, C., Besnard, A.-L., Launey, S., Evanno, G., 2017. Inferring the demographic history underlying parallel genomic divergence among pairs of parasitic and nonparasitic lamprey ecotypes. Mol. Ecol. 26, 142–162. 10.1111/mec.13664

69. Rougeux, C., Gagnaire, P.-A., Bernatchez, L., 2019. Model-based demographic inference of introgression history in European whitefish species pairs’. J. Evol. Biol. 32, 806–817. 10.1111/jeb.13482

70. Saltin, S.H., Schade, H., Johannesson, K., 2013. Preference of males for large females causes a partial mating barrier between a large and a small ecotype of Littorina fabalis (W. Turton, 1825). J. Molluscan Stud. 79, 128–132. 10.1093/mollus/eyt003

71. Schliep, K.P., 2011. phangorn: phylogenetic analysis in R. Bioinformatics 27, 592–593. 10.1093/bioinformatics/btq706

72. Silva, A.C.F., Hawkins, S., Boaventura, D., Brewster, E., Thompson, R.C., 2010. Use of the intertidal zone by mobile predators: Influence of wave exposure, tidal phase and elevation on abundance and diet. Mar. Ecol. Prog. Ser. 406, 197–210. 10.3354/meps08543

73. Soria-Carrasco, V., Gompert, Z., Comeault, A.A., Farkas, T.E., Parchman, T.L., Johnston, J.S., Buerkle, C.A., Feder, J.L., Bast, J., Schwander, T., Egan, S.P., Crespi, B.J., Nosil, P., 2014. Stick insect genomes reveal natural selection’s role in parallel speciation. Science 344, 738–742. 10.1126/science.1252136

74. Sotelo, G., Duvetorp, M., Costa, D., Panova, M., Johannesson, K., Faria, R., 2020. Phylogeographic history of flat periwinkles, Littorina fabalis and L. obtusata. BMC Evol. Biol. 20, 23. 10.1186/s12862-019-1561-6

75. Tan, T., Atkinson, E.G., 2023. Strategies for the Genomic Analysis of Admixed Populations. Annu. Rev. Biomed. Data Sci. 6, 105–127. 10.1146/annurev-biodatasci-020722-014310

76. Tatarenkov, A., Johannesson, K., 1999. Micro- and macrogeographic allozyme variation in Littorina fabalis ; do sheltered and exposed forms hybridize? Biol. J. Linn. Soc. 67, 199–212. 10.1006/bijl.1998.0303

77. Tatarenkov, A., Johannesson, K., 1998. Evidence of a reproductive barrier between two forms of the marine periwinkle Littorina fabalis (Gastropoda). Biol. J. Linn. Soc. 63, 349–365. 10.1111/j.1095-8312.1998.tb01522.x

78. Tatarenkov, A., Johannesson, K., 1994. Habitat related allozyme variation on a microgeographic scale in the marine snail Littorina mariae (Prosobranchia: Littorinacea). Biol. J. Linn. Soc. 53, 105–125. 10.1111/j.1095-8312.1994.tb01004.x

79. Weir, B.S., Cockerham, C.C., 1984. Estimating F-Statistics for the Analysis of Population Structure. Evolution 38, 1358–1370. 10.2307/2408641

80. Wellenreuther, M., Bernatchez, L., 2018. Eco-Evolutionary Genomics of Chromosomal Inversions. Trends Ecol. Evol. 33, 427–440. 10.1016/j.tree.2018.04.002

81. Westram, A.M., Faria, R., Johannesson, K., Butlin, R., 2021. Using replicate hybrid zones to understand the genomic basis of adaptive divergence. Mol. Ecol. 30, 3797–3814. 10.1111/mec.15861

82. Westram, A.M., Faria, R., Johannesson, K., Butlin, R., Barton, N., 2022. Inversions and parallel evolution. Philos. Trans. R. Soc. B Biol. Sci. 377, 20210203. 10.1098/rstb.2021.0203

83. Yang, J., Benyamin, B., McEvoy, B.P., Gordon, S., Henders, A.K., Nyholt, D.R., Madden, P.A., Heath, A.C., Martin, N.G., Montgomery, G.W., Goddard, M.E., Visscher, P.M., 2010. Common SNPs explain a large proportion of heritability for human height. Nat. Genet. 42, 565–569. 10.1038/ng.608

84. Yu, G., Lam, T.T.-Y., Zhu, H., Guan, Y., 2018. Two Methods for Mapping and Visualizing Associated Data on Phylogeny Using Ggtree. Mol. Biol. Evol. 35, 3041–3043. 10.1093/molbev/msy194

