## Supplementary Material for "Inversions support both parallel and location-specific adaptations in snail ecotypes"

### Supplementary Materiel

Table 1: Distribution parameters of shell size cline and brown colour frequency cline. Mean shell size values in the sheltered and exposed parts are indicated in millimeters, and the colour values are expressed as percent brown individuals. The centers and widths of the clines are indicated in meters. The values in square brackets represent the 95 % confidence intervals on the estimated parameters. The ΔAIC is the difference between the best model's AIC and the second-best model's AIC (note that the stable and linear models could not be decided between)

| Trait | Country | Phenotypic trait value | | Cline center (m) | Cline width (m) | Best model | ΔAIC |
|---|---|---|---|---|---|---|---|
| | | Sheltered part | Exposed part | | | | |
| Size | Sweden | 8.2 [7.1, 8.9] | 11.5 [11.2, 11.9] | 64.3 [47.6, 75.0] | 51.1 [17.4, 112.9] | Clinal | / |
| | France | 10.9 [10.6, 11.2] | 8.4 [7.9, 8.8] | 217.2 [205.5, 237.4] | 46.1 [21.5, 93.2] | Clinal | / |
| Color | Sweden | 7.7% [4.3, 12.4] | 7.7% [4.3, 12.4] | / | / | Stable | -0.7. |
| | France | 36.1% [18.7, 48.4] | 90.9% [74.0, 100.0] | 235.0 [208.9, 275.4] | 56.4 [0.4, 212.1] | Clinal | -7.5 |

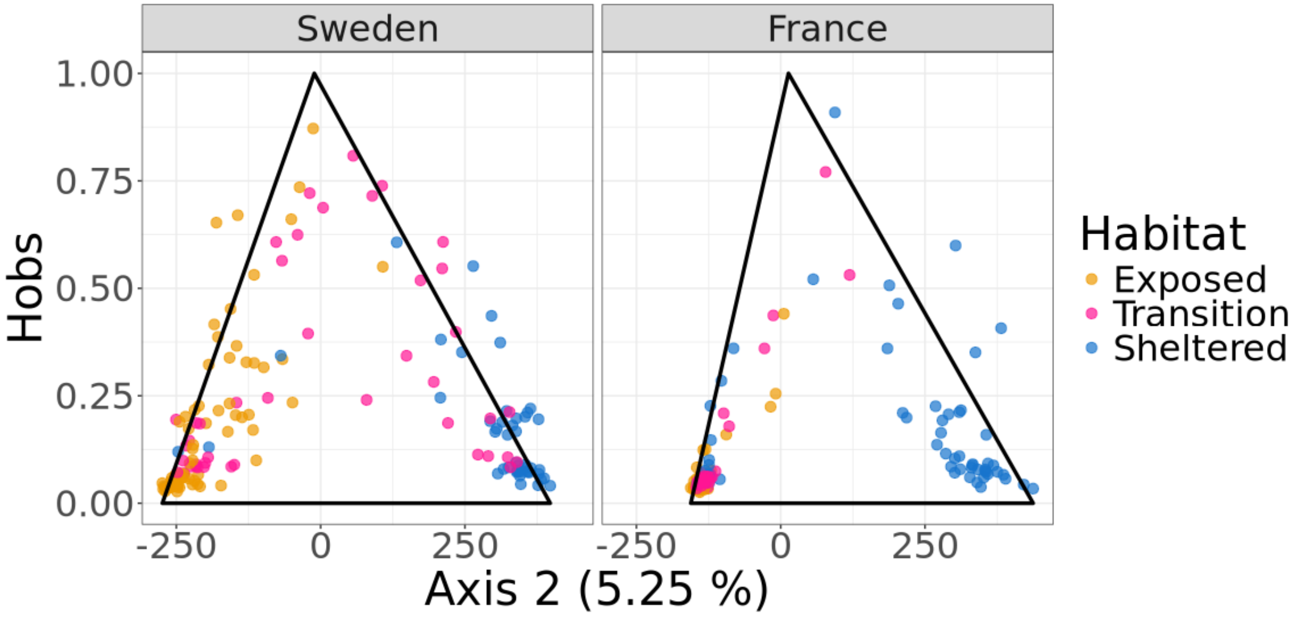


**Figure S2:** Correlation between individual scores on the second principal component (PC2) and observed heterozygosity (Hobs) calculated from SNPs with Δfreq > 0.80 between ecotypes. Individuals are colored according to their habitat. Snails with intermediate positions on PC2 (associated with ecotype differentiation in both France and Sweden) exhibit high levels of heterozygosity at SNPs that are nearly diagnostic of ecotype, consistent with these individuals being recent hybrids between sheltered and exposed ecotypes.

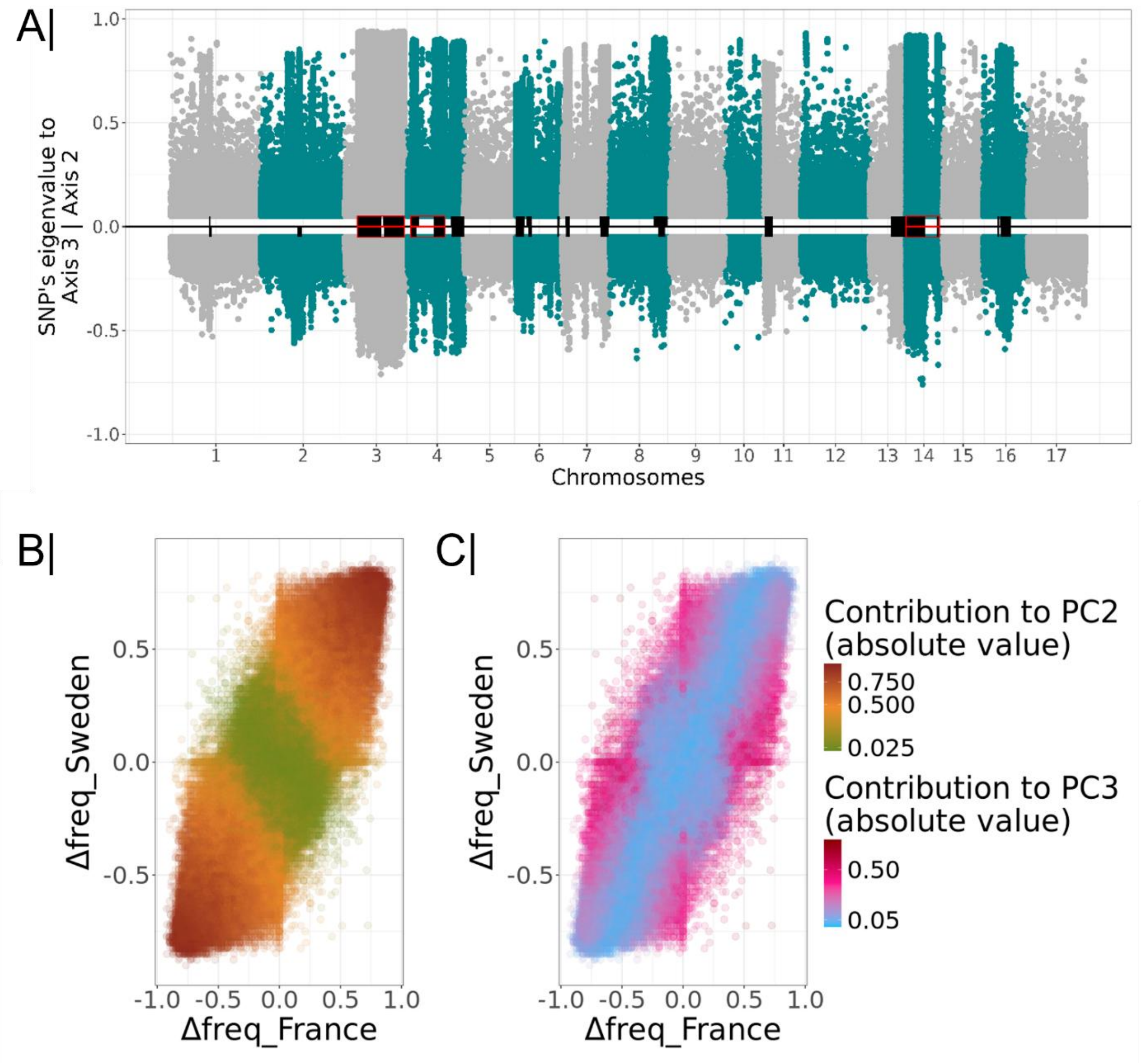


**Figure S3:** Summary of SNP contributions (eigenvalues) to individual dispersion in the PCA and to ecotype differentiation. **(A)** Distribution of SNP contributions to PC2 (above 0) and PC3 (below 0) from the PCA analyses presented in Figure 1 of the main text. **(B)** and **(C)** Correlation between inter-ecotype allele frequency differences (Δfreq) for each SNP in Sweden and France. Points are colored according to the absolute value of each SNP's contribution to **(B)** PC2 (parallel associations between genetic variation and the wave exposure gradient in Sweden and France) or **(C)** PC3 (reverse associations between genetic variation and the wave exposure gradient in Sweden and France). The vast majority of SNPs with strong contributions to PC2 (brown points) show large Δfreq values (i.e., highly negative or highly positive) that are similar between the two study sites. Most SNPs whose genetic variation is captured by PC3 (reverse association with the wave exposure gradient) have high Δfreq in only one country (red points forming a "cross" pattern, panel C). Overall, these results show that the genomic regions contributing to the dispersion of snails on PC2 and PC3 (the two PCs correlated with the environmental gradient) are broadly the same **(A)**, but that the specific SNPs contributing to the dispersion differ between the two axes (B & C).

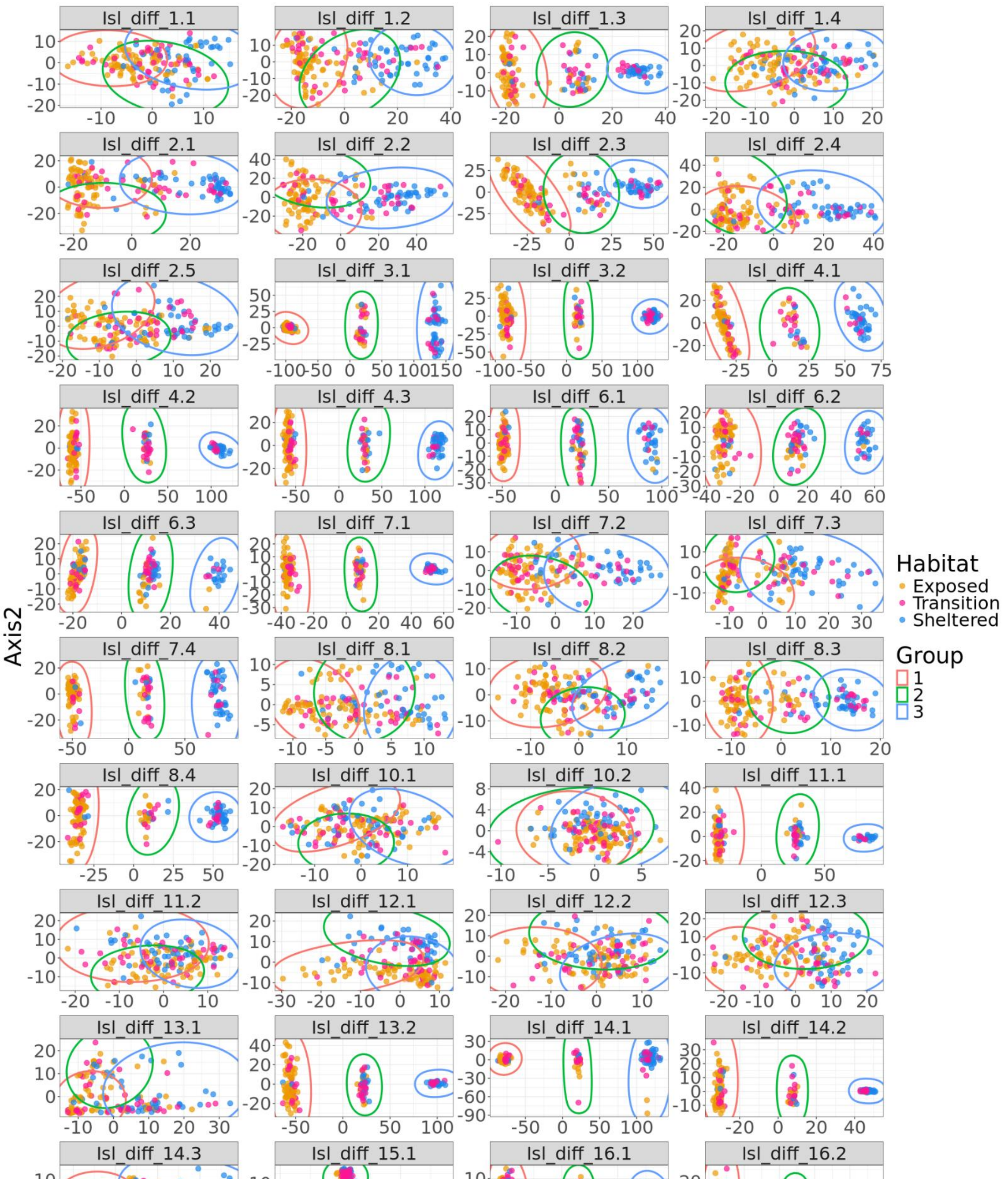


Figure S4: Local PCA on islands of differentiation detected by the HMM in Sweden. Each facet represents a different island of differentiation. Each point represents an individual and the points are coloured depending on the individual's habitat (Exposed in orange, Transition in pink and Sheltered in blue). The ellipses represent the results of the DAPC approach. Although the signal of high differentiation on LG15 (LG15.1) is clearly showing three discrete clusters, these clusters are linked to sex determining locus, and were kept out of our analyses as it is out of the scope of our manuscript.

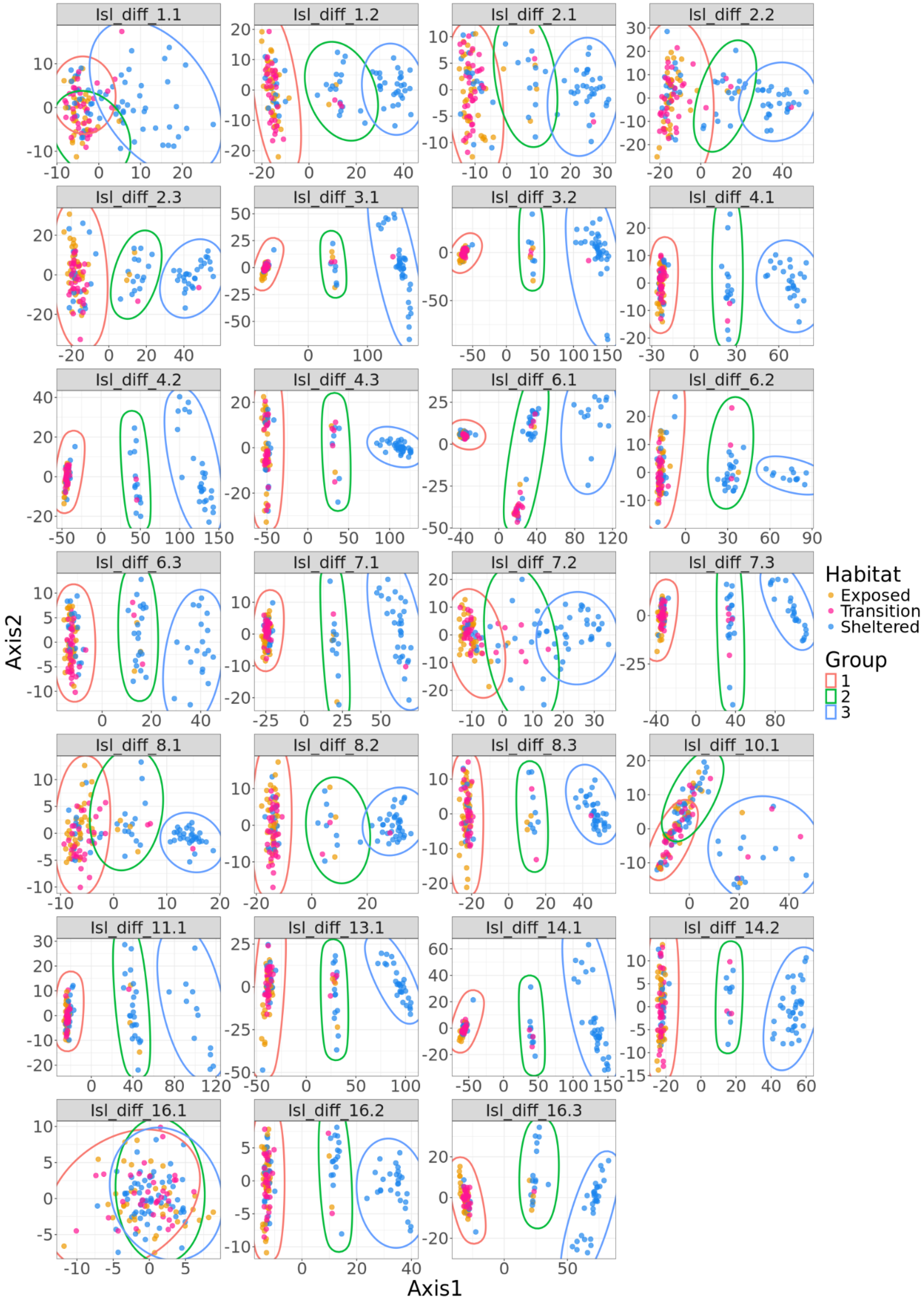


Figure S5: Local PCA on islands of differentiation detected by the HMM in France. Each facet represents a different island of differentiation. Each point represents an individual and the points are coloured depending on the individual's habitat (Exposed in orange, Transition in pink and Sheltered in blue). The ellipses represent the results of the DAPC approach.

**Figure S6:** Individual heterozygosity for each of the three distinct clusters detected within the genomic island matching criterion iii (presence of three distinct clusters with one cluster located at mid-distance between the two others) in our inversion detection analyses in Sweden (two top rows) and France (two bottom rows). The clusters are labeled E/E, E/S, and S/S, corresponding to the three potential karyotypes for each inversion. All graphs show that samples in the central cluster (labeled E/S) have much higher heterozygosity than samples from the two outer clusters of the local PCA in Figures S4–S5, as expected for individuals that are heterozygous for an inversion.

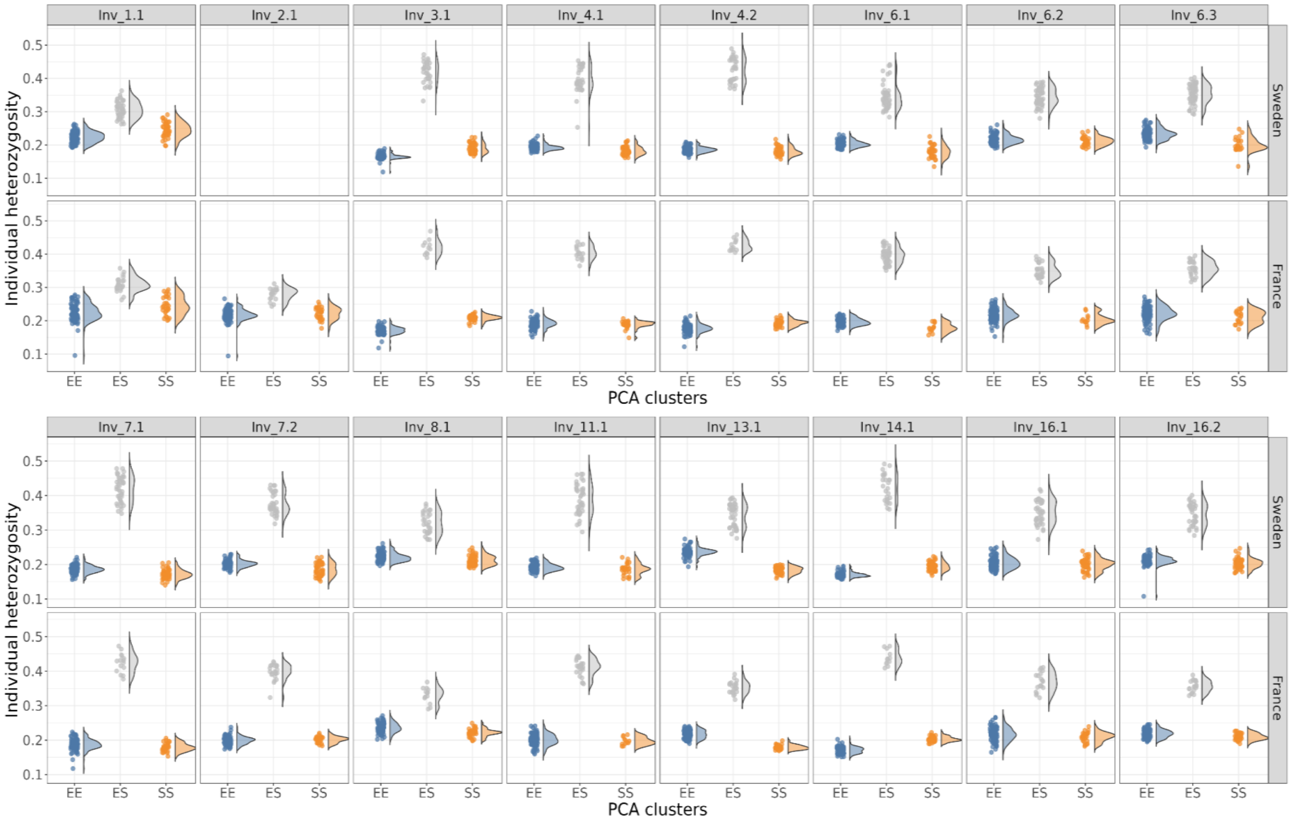

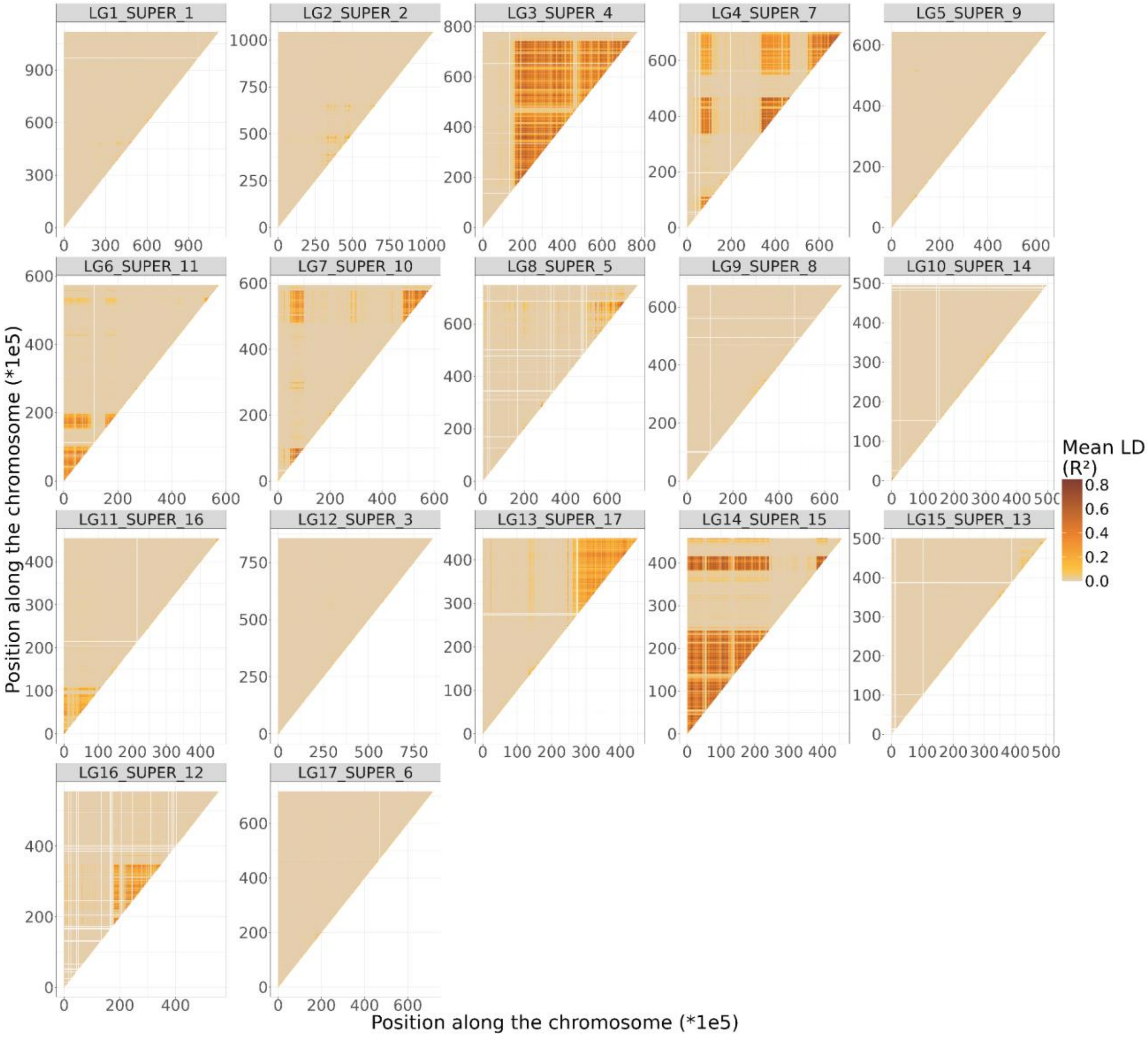


Figure S7: Mean linkage disequilibrium per 10kb bin for each chromosome in Sweden.

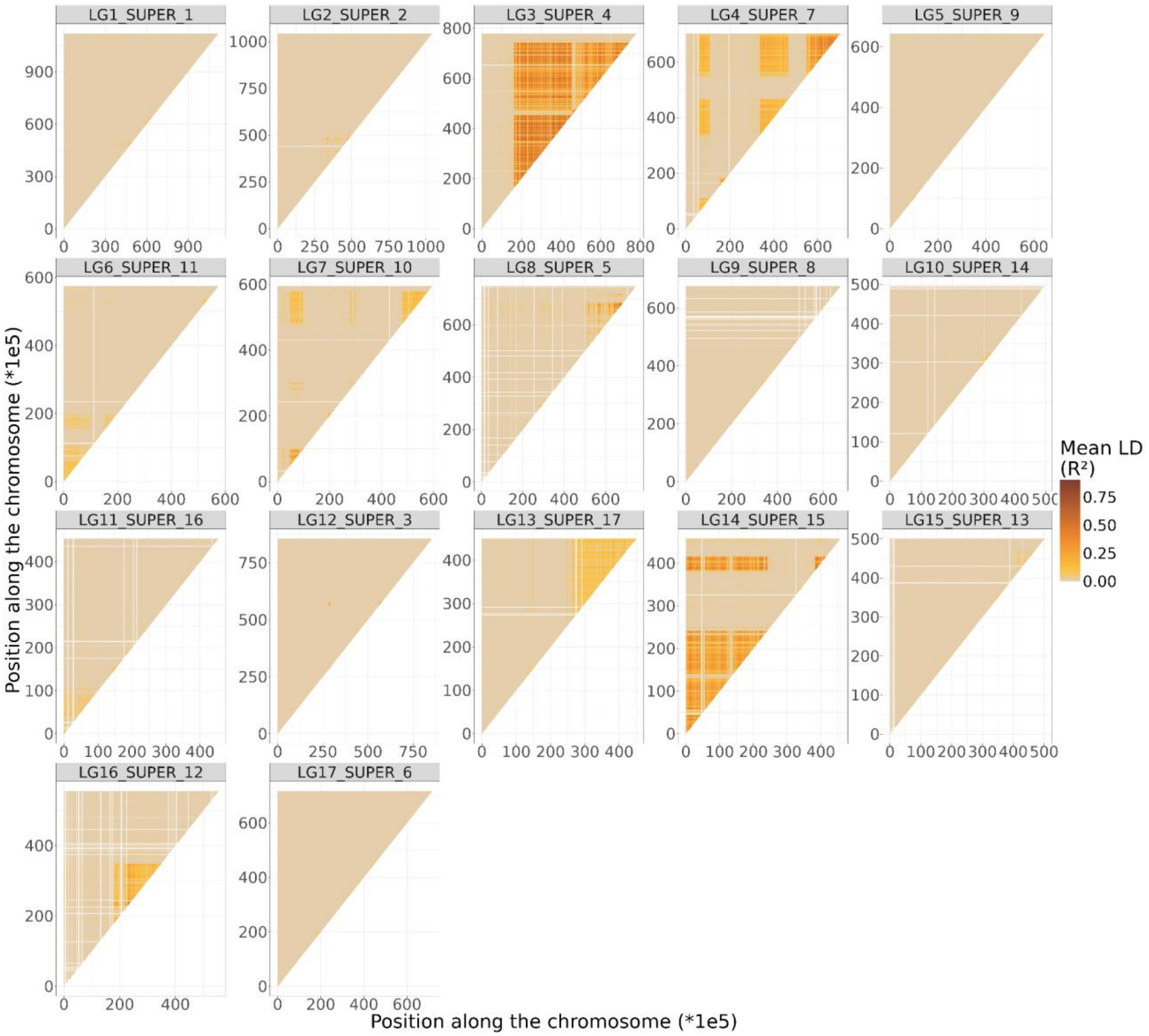


Figure S8: Mean linkage disequilibrium per 10kb bin for each chromosome in France

**Table S2:** Population statistics within chromosomal inversions identified by local PCA in Sweden and France using the overall dataset (507k SNPs thinned for physical linkage disequilibrium). LD was computed over all French samples or over all Swedish samples. Heterozygosity was computed for each potential karyotype of the inversions, labeled EE/ES/SS, corresponding to the observed heterozygosity within each inversion genotype; "–" indicates missing data for inversions present in one country but absent in the other.

| | **Sweden** | | | | **France** | | | |
|---|---|---|---|---|---|---|---|---|
| **Inversion** | LD | EE | ES | SS | LD | EE | ES | SS |
| Inv_1.1 | 0.07 | 0.23 | 0.31 | 0.25 | 0.04 | 0.22 | 0.30 | 0.24 |
| Inv_2.1 | - | - | - | - | 0.03 | 0.22 | 0.28 | 0.22 |
| Inv_3.1 | 0.40 | 0.16 | 0.44 | 0.22 | 0.40 | 0.15 | 0.41 | 0.18 |
| Inv_3.2 | 0.37 | 0.19 | 0.41 | 0.19 | 0.30 | 0.18 | 0.38 | 0.18 |
| Inv_4.1 | 0.16 | 0.22 | 0.34 | 0.21 | 0.05 | 0.21 | 0.33 | 0.21 |
| Inv_4.2 | 0.43 | 0.18 | 0.45 | 0.17 | 0.15 | 0.18 | 0.42 | 0.17 |
| Inv_4.3 | 0.33 | 0.18 | 0.43 | 0.19 | 0.27 | 0.19 | 0.40 | 0.18 |
| Inv_6.1 | 0.26 | 0.20 | 0.40 | 0.18 | 0.07 | 0.20 | 0.34 | 0.19 |
| Inv_6.2 | 0.14 | 0.22 | 0.35 | 0.21 | 0.03 | 0.22 | 0.33 | 0.21 |
| Inv_6.3 | 0.11 | 0.22 | 0.36 | 0.21 | 0.05 | 0.23 | 0.34 | 0.20 |
| Inv_7.1 | 0.39 | 0.19 | 0.43 | 0.18 | 0.20 | 0.19 | 0.40 | 0.17 |
| Inv_7.2 | 0.28 | 0.20 | 0.40 | 0.20 | 0.09 | 0.20 | 0.36 | 0.18 |
| Inv_8.1 | 0.13 | 0.24 | 0.33 | 0.22 | 0.10 | 0.22 | 0.30 | 0.21 |
| Inv_11.1 | 0.14 | 0.20 | 0.42 | 0.20 | 0.04 | 0.19 | 0.36 | 0.18 |
| Inv_13.1 | 0.24 | 0.22 | 0.36 | 0.18 | 0.09 | 0.24 | 0.32 | 0.19 |
| Inv_14.1 | 0.43 | 0.17 | 0.44 | 0.20 | 0.23 | 0.17 | 0.39 | 0.20 |
| Inv_14.2 | 0.51 | 0.18 | 0.47 | 0.18 | 0.31 | 0.16 | 0.41 | 0.17 |
| Inv_16.1 | 0.21 | 0.22 | 0.37 | 0.21 | 0.08 | 0.20 | 0.21 | 0.25 |
| Inv_16.2 | 0.24 | 0.22 | 0.36 | 0.21 | 0.14 | 0.21 | 0.33 | 0.20 |

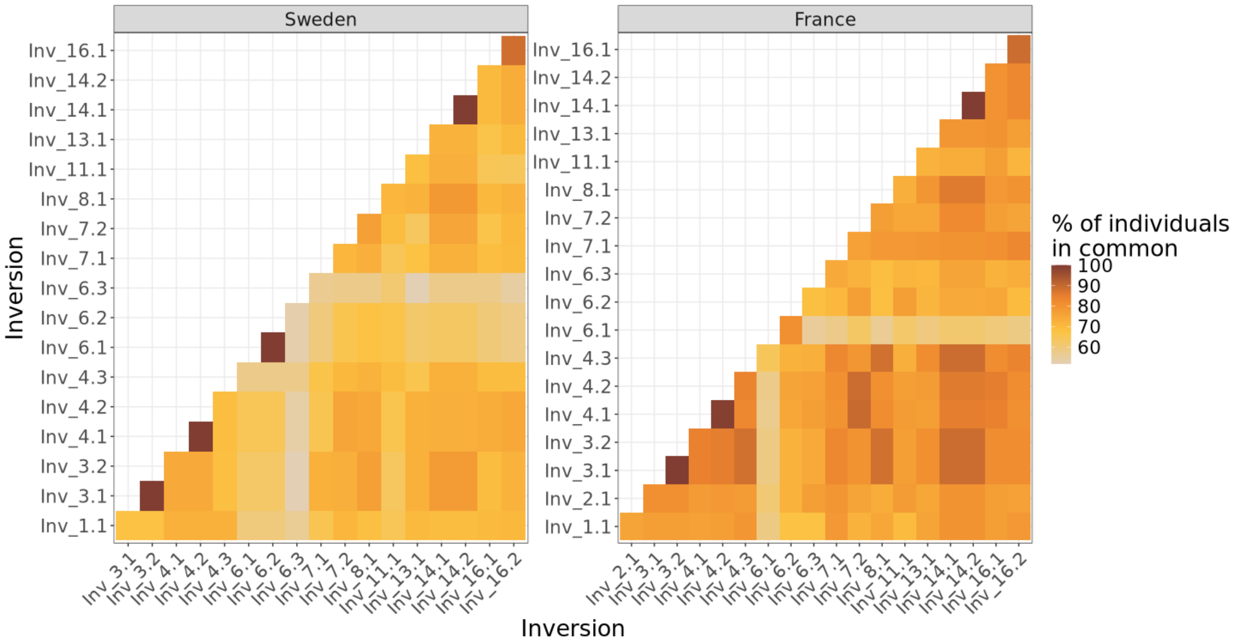


**Figure S9:** Percentage of individuals sharing the same PCA cluster across different inversions detected using our 4-criterium approach. This analysis shows that 4 pairs of inversions in Sweden and three pairs of inversion in France are considered separated based on our four-criterion approach but are in complete linkage disequilibrium (LD=1). For example, for Inv_3.1 and Inv_3.2 in both France and Sweden, 100% of individuals with a given karyotype for Inv_3.1 also have the same karyotype for Inv_3.2. Although we acknowledge that more complex rearrangements involving multiple inversions and partial recollinearization of a portion of the chromosome could also produce this pattern of high LD between distant chromosomal regions, we use a parsimonious argument to consider each of these pairs of putative inversions in complete LD as a single inversion whose apparent split is an artifact of using a sister species as a reference genome.

Table S3: Identified putative inversions and breakpoints. The values indicated are in units of 100kbp. Inversions with similar breakpoints in Sweden and France. The numbers with bars (ex. 153/478) represent the portions of putative inversions that were grouped together with strict linkage disequilibrium. The calculated length of the inversion is the sum of the two portions if there are two.

| Chromosome | Inversion | Sweden | | | France | | |
|---|---|---|---|---|---|---|---|
| | | Start | End | Length | Start | End | Length |
| LG1 | Inv_1.1 | 475 | 492 | 17 | 475 | 507 | 32 |
| LG2 | Inv_2.1 | / | / | / | 452 | 510 | 58 |
| LG3 | Inv_3.1 | 153/478 | 454/742 | 565 | 155/488 | 453/742 | 552 |
| LG4 | Inv_4.1 | 43/330 | 146/446 | 239 | 43/330 | 109/466 | 202 |
| LG4 | Inv_4.2 | 538 | 702 | 164 | 549 | 702 | 153 |
| LG6 | Inv_6.1 | 1 | 112 | 111 | 1 | 104 | 103 |
| LG6 | Inv_6.2 | 137 | 196 | 56 | 155 | 197 | 42 |
| LG6 | Inv_6.3 | 518 | 542 | 24 | 518 | 536 | 18 |
| LG7 | Inv_7.1 | 41 | 98 | 57 | 47 | 98 | 51 |
| LG7 | Inv_7.2 | 470 | 588 | 118 | 478 | 577 | 99 |
| LG8 | Inv_8.1 | 540 | 717 | 177 | 600 | 684 | 84 |
| LG11 | Inv_11.1 | 0 | 107 | 107 | 0 | 106 | 106 |
| LG13 | Inv_13.1 | 260 | 449 | 189 | 265 | 445 | 180 |
| LG14 | Inv_14.1 | 1/382 | 241/415 | 273 | 1/380 | 241/415 | 275 |
| LG16 | Inv_16.1 | 180 | 203 | 23 | 180 | 196 | 16 |
| LG16 | Inv_16.2 | 213 | 348 | 135 | 216 | 348 | 132 |

**Table S4:** Pairwise $F_{ST}$ matrices computed using only the references samples, and the SNPs associated with ecotype differences in Sweden only. The label FR stand for the samples from France, SW for Sweden, _S for sheltered habitat and _E for exposed habitat.

| | **FR_E** | **FR_S** | **SW_S** |
|---|---|---|---|
| **FR_S** | 0.031 | | |
| **SW_S** | 0.554 | 0.489 | |
| **SW_E** | 0.235 | 0.276 | 0.612 |

**Table S5:** Pairwise $F_{ST}$ matrices computed using only the references samples, and the SNPs associated with ecotype differences in France only. The label FR stand for the samples from France, SW for Sweden, _S for sheltered habitat and _E for exposed habitat.

| | **FR_E** | **FR_S** | **SW_S** |
|---|---|---|---|
| **FR_S** | 0.587 | | |
| **SW_S** | 0.531 | 0.398 | |
| **SW_E** | 0.500 | 0.444 | 0.030 |

**Table S6:** Pairwise $F_{ST}$ matrices computed using only the references samples, and the SNPs associated with ecotype differences in both country but whose allele are reversed. The label FR stand for the samples from France, SW for Sweden, _S for sheltered habitat and _E for exposed habitat.

| | **FR_E** | **FR_S** | **SW_S** |
|---|---|---|---|
| **FR_S** | 0.483 | | |
| **SW_S** | 0.208 | 0.382 | |
| **SW_E** | 0.550 | 0.119 | 0.401 |

**Legend:** FR_E = France Exposed, FR_S = France Sheltered, SW_S = Sweden Sheltered, SW_E = Sweden Exposed

**Table S5:** Parameters of frequency clines inferred for putative chromosomal inversions. Frequencies are in percentages. Values shown as mean [2.5%, 97.5%].

| Pop | Inv | Centre (m) | Width (m) | Left | Right | $F_{IS}$ |
|---|---|---|---|---|---|---|
| Sweden | Inv_1.1 | 65.23 [55.40, 70.81] | 40.88 [20.89, 76.23] | 0.86 [0.75, 0.97] | 0.12 [0.07, 0.18] | 0.74 [0.32, 0.99] |
| | Inv_3.1 | 67.94 [61.13, 74.71] | 56.46 [38.28, 79.35] | 0.99 [0.95, 1.00] | 0.08 [0.03, 0.15] | 0.51 [0.19, 0.78] |
| | Inv_4.1 | 60.67 [52.13, 68.91] | 55.69 [31.89, 80.99] | 0.97 [0.85, 1.00] | 0.06 [0.02, 0.11] | 0.61 [0.27, 0.91] |
| | Inv_4.2 | 59.37 [50.25, 68.09] | 75.97 [40.40, 112.62] | 0.97 [0.84, 1.00] | 0.03 [0.00, 0.08] | 0.55 [0.21, 0.85] |
| | Inv_6.1 | 56.41 [36.78, 79.45] | 90.95 [26.58, 119.62] | 0.88 [0.63, 1.00] | 0.10 [0.01, 0.19] | 0.23 [-0.11, 0.57] |
| | Inv_6.2 | 56.77 [35.56, 78.88] | 91.48 [25.30, 119.43] | 0.88 [0.62, 1.00] | 0.10 [0.02, 0.18] | 0.23 [-0.07, 0.52] |
| | Inv_6.3 | 73.89 [57.54, 81.14] | 25.46 [12.20, 67.08] | 0.57 [0.46, 0.73] | 0.12 [0.08, 0.18] | -0.46 [-0.97, 0.17] |
| | Inv_7.1 | 61.95 [52.59, 70.78] | 67.11 [31.59, 106.43] | 0.97 [0.84, 1.00] | 0.11 [0.05, 0.18] | 0.58 [0.26, 0.89] |
| | Inv_7.2 | 61.18 [51.81, 69.95] | 65.03 [42.42, 92.65] | 0.99 [0.92, 1.00] | 0.08 [0.02, 0.15] | 0.51 [0.16, 0.84] |
| | Inv_8.1 | 65.97 [59.38, 72.12] | 60.29 [40.23, 88.30] | 0.99 [0.96, 1.00] | 0.05 [0.00, 0.11] | 0.70 [0.32, 0.97] |
| | Inv_11.1 | 62.01 [42.65, 75.28] | 66.79 [19.76, 118.64] | 0.77 [0.58, 1.00] | 0.03 [0.00, 0.10] | 0.38 [-0.05, 0.76] |
| | Inv_13.1 | 65.47 [55.08, 74.90] | 75.77 [32.47, 114.37] | 0.92 [0.76, 1.00] | 0.04 [0.00, 0.12] | 0.38 [0.01, 0.74] |
| | Inv_14.1 | 69.94 [61.55, 77.72] | 79.54 [60.31, 102.07] | 0.99 [0.96, 1.00] | 0.01 [0.00, 0.05] | 0.62 [0.34, 0.85] |
| | Inv_16.1 | 61.42 [55.02, 69.12] | 56.56 [33.88, 79.19] | 0.98 [0.88, 1.00] | 0.10 [0.05, 0.16] | 0.52 [0.16, 0.84] |
| | Inv_16.2 | 64.35 [57.28, 71.44] | 42.83 [25.21, 63.02] | 0.99 [0.91, 1.00] | 0.10 [0.05, 0.16] | 0.50 [0.10, 0.88] |
| France | Inv_1.1 | 160.71 [136.59, 176.99] | 103.75 [50.04, 166.67] | 0.93 [0.81, 1.00] | 0.01 [0.00, 0.06] | 0.48 [-0.01, 0.94] |
| | Inv_2.1 | 178.52 [151.13, 194.02] | 116.40 [61.67, 188.24] | 0.73 [0.57, 0.89] | 0.01 [0.00, 0.04] | 0.80 [0.52, 0.98] |
| | Inv_3.1 | 176.56 [153.98, 190.29] | 130.86 [78.56, 217.19] | 0.88 [0.75, 1.00] | 0.01 [0.00, 0.03] | 0.94 [0.79, 1.00] |
| | Inv_4.1 | 166.57 [125.02, 182.97] | 93.50 [55.67, 186.13] | 0.81 [0.66, 1.00] | 0.00 [0.00, 0.02] | 0.74 [0.40, 0.97] |
| | Inv_4.2 | 160.75 [130.80, 180.27] | 118.22 [56.99, 203.05] | 0.95 [0.85, 1.00] | 0.01 [0.00, 0.05] | 0.80 [0.47, 0.99] |
| | Inv_6.1 | 199.31 [158.05, 240.97] | 161.55 [70.94, 225.96] | 0.56 [0.38, 0.76] | 0.01 [0.00, 0.08] | -0.13 [-0.40, 0.13] |
| | Inv_6.2 | 151.47 [81.52, 196.51] | 143.51 [58.16, 226.51] | 0.61 [0.38, 1.00] | 0.01 [0.00, 0.04] | 0.31 [-0.07, 0.71] |
| | Inv_6.3 | 200.07 [172.39, 215.47] | 114.89 [58.15, 202.12] | 0.51 [0.38, 0.67] | 0.01 [0.00, 0.05] | 0.78 [0.42, 0.99] |
| | Inv_7.1 | 170.83 [141.58, 183.23] | 86.23 [41.57, 168.57] | 0.86 [0.73, 1.00] | 0.01 [0.00, 0.05] | 0.80 [0.46, 0.99] |
| | Inv_7.2 | 170.89 [148.61, 184.98] | 89.17 [52.28, 143.48] | 0.77 [0.62, 0.90] | 0.00 [0.00, 0.02] | 0.64 [0.28, 0.92] |
| | Inv_8.1 | 173.17 [147.34, 191.91] | 121.66 [67.20, 201.58] | 0.93 [0.80, 1.00] | 0.01 [0.00, 0.04] | 0.88 [0.65, 0.99] |
| | Inv_11.1 | 148.93 [73.70, 184.51] | 103.58 [22.24, 227.16] | 0.65 [0.40, 1.00] | 0.02 [0.00, 0.06] | 0.54 [-0.02, 0.98] |
| | Inv_13.1 | 148.17 [120.04, 171.94] | 124.58 [27.91, 209.88] | 0.95 [0.82, 1.00] | 0.03 [0.00, 0.10] | 0.69 [0.26, 0.96] |
| | Inv_14.1 | 179.53 [162.61, 188.83] | 71.81 [46.99, 110.21] | 0.85 [0.73, 0.95] | 0.00 [0.00, 0.02] | 0.85 [0.57, 1.00] |
| | Inv_16.1 | 154.85 [116.94, 176.75] | 121.67 [60.45, 213.91] | 0.90 [0.75, 1.00] | 0.01 [0.00, 0.04] | 0.57 [0.17, 0.88] |
| | Inv_16.2 | 168.65 [129.19, 187.83] | 88.56 [42.59, 200.74] | 0.86 [0.72, 1.00] | 0.03 [0.00, 0.08] | 0.63 [0.25, 0.94] |

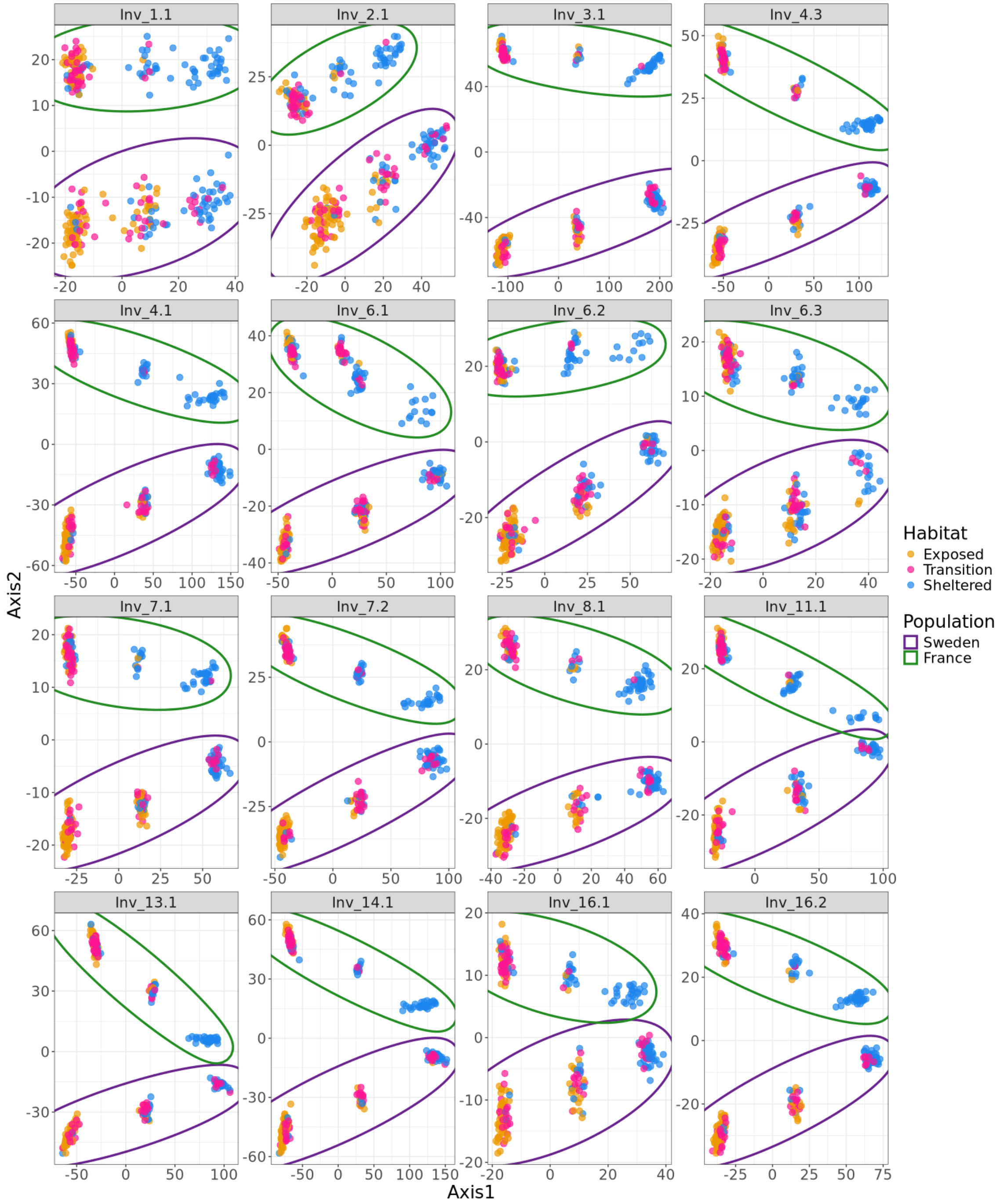


Figure S10: Local PCA analyses on islands of differentiation both in Sweden and France. Each graph represents PC1 against PC2. We ran the local PCA in both populations only in the regions identified as inversions using the local PCAs on the islands of differentiation in the separated populations (Fig. S5-S6). The names of the inversions and their genomic boundaries correspond to the ones indicated in Table S3.

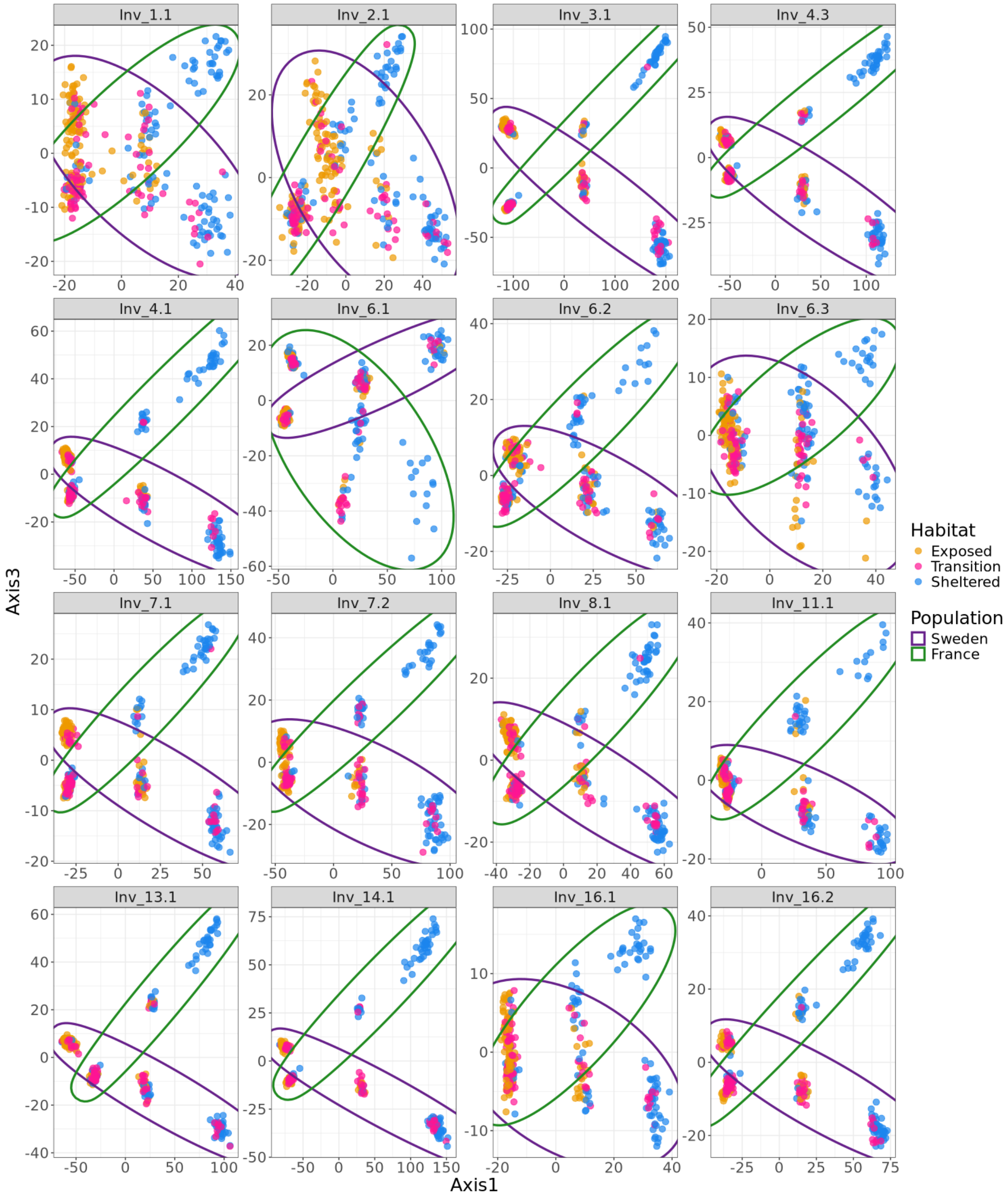


Figure S11: Local PCA analyses on islands of differentiation both in Sweden and France. Each graph represents PC1 against PC3. We ran the local PCA in both populations only in the regions identified as inversions using the local PCAs on the islands of differentiation in the separated populations (Fig. S5-S6). The names of the inversions and their genomic boundaries correspond to the ones indicated in Table S3.

Table S6: Divergence between karyotypes of the inversion. Only the homokaryotype individuals were used here. The decimal values are indicated in percentages. The blue individuals in the trees are the Swedish individuals. The light blue individuals are the individuals with the SS karyotype, the dark blue individuals are the individuals with the EE karyotype. The individuals in red and orange are the French individuals. The orange ones are the SS karyotype and the red ones are the EE karyotype. The parts of the genome that have names starting with “Inv” are inversions and the ones starting with “Col” are colinear genome. The coordinates of these portions can be found in the following table.

| Portion of genome | Tree | Divergence between karyotypes | |
|---|---|---|---|
| | | Sweden | France |
| Inv_1.1 | 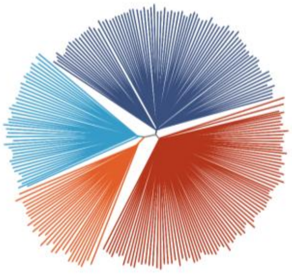 | 31.8 | 33.3 |
| Inv_2.1 | 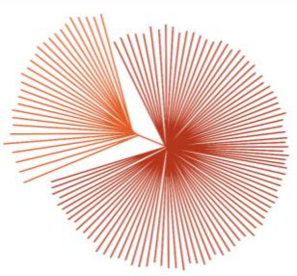 | / | 38.7 |
| Inv_3.1 | 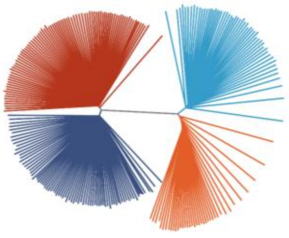 | 48.0 | 48.6 |
| Inv_4.1 | 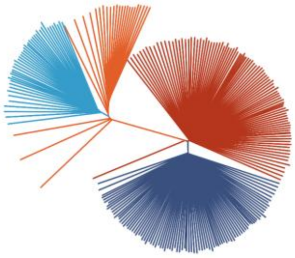 | 50.4 | 49.7 |

| | | | |
|---|---|---|---|
| Inv_4.2 | 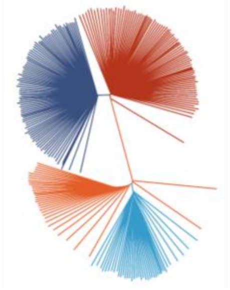 | 44.8 | 44.5 |
| Inv_6.1 | 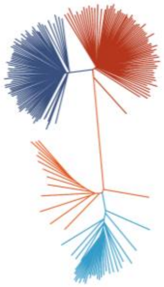 | 57.3 | 50.3 |
| Inv_6.2 | 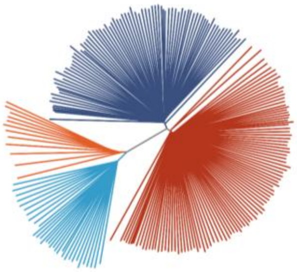 | 39.6 | 40.5 |
| Inv_6.3 | 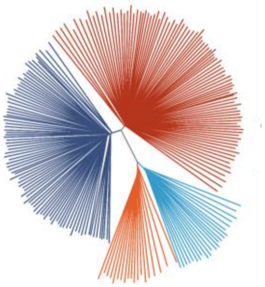 | 36.9 | 37.8 |
| Inv_7.1 | 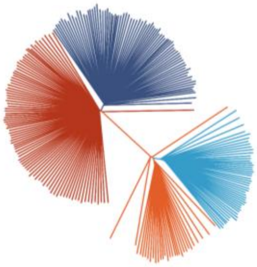 | 53.6 | 52.5 |

| | | | |
|---|---|---|---|
| Inv_7.2 | 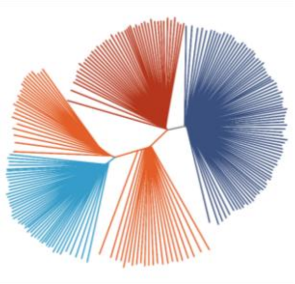 | 46.4 | 42.5 |
| Inv_8.1 | 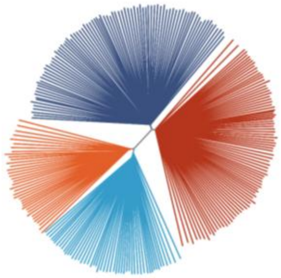 | 44.2 | 44.1 |
| Inv_11.1 | 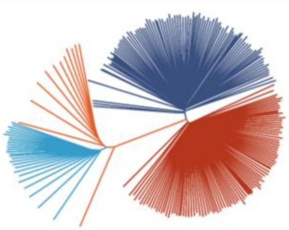 | 44.0 | 43.4 |
| Inv_13.1 | 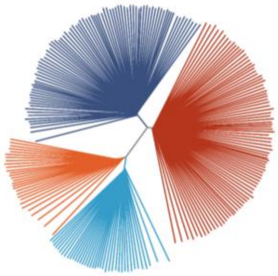 | 42.9 | 42.7 |
| Inv_14.1 | 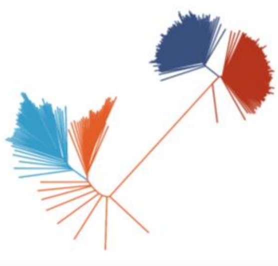 | 68.7 | 59.6 |

| | | | |
|---|---|---|---|
| Inv_16.1 | 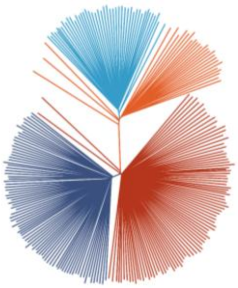 | 43.6 | 42.3 |
| Inv_16.2 | 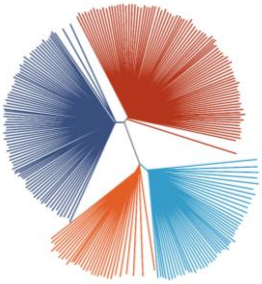 | 38.6 | 38.6 |
| Col_1.1 | 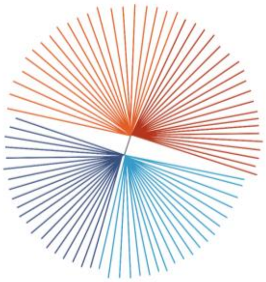 | 36.8 | 39.4 |
| Col_3.1 | 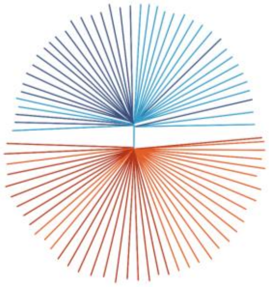 | 36.7 | 40.3 |
| Col_4.1 | 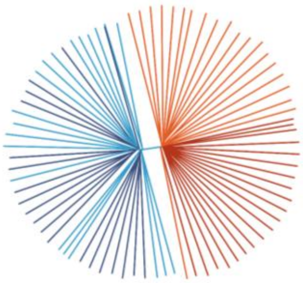 | 34.8 | 37.2 |

| | | | |
|---|---|---|---|
| Col_5.1 | 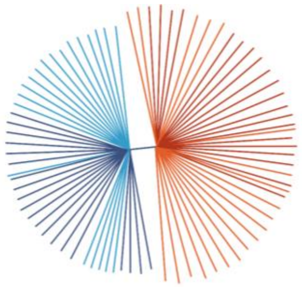 | 35.2 | 39.5 |
| Col_6.1 | 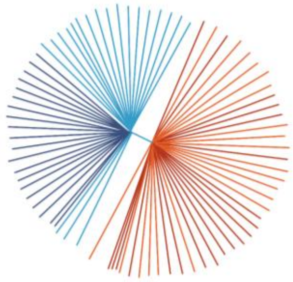 | 35.8 | 38.5 |
| Col_11.1 | 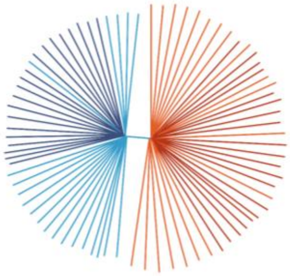 | 35.5 | 38.8 |
| Col_14.1 | 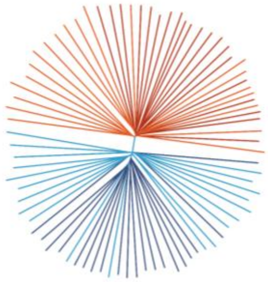 | 39.5 | 41.1 |
| Col_17.1 | 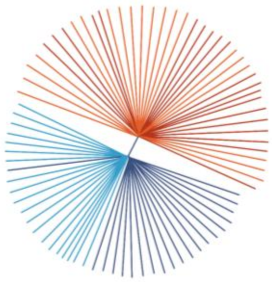 | 36.1 | 38.8 |

*<u>Table S7</u>: Delimitation of the collinear parts of the genome used to represent control trees. The values indicated are in units of 1e5 bp.*

| Chromosome | Collinear part | France & Sweden | | |
|---|---|---|---|---|
| | | Start | End | Length |
| LG1 | Col_1.1 | 0 | 475 | 475 |
| LG3 | Col_3.1 | 0 | 153 | 153 |
| LG4 | Col_4.1 | 146 | 330 | 184 |
| LG5 | Col_5.1 | 100 | 600 | 500 |
| LG6 | Col_6.1 | 200 | 510 | 310 |
| LG11 | Col_11.1 | 110 | 450 | 340 |
| LG14 | Col_14.1 | 415 | 457 | 42 |
| LG17 | Col_17.1 | 0 | 72 | 72 |
